# Molecular and functional characterization of buffalo nasal epithelial odorant binding proteins and their structural insights by *in-silico* and biochemical approach

**DOI:** 10.1101/2020.09.17.301234

**Authors:** Chidhambaram Manikkaraja, Bhavika Mam, Randhir Singh, Balasubramanian Nagarathnam, Geen George, Akash Gulyani, Govindharaju Archunan, Ramanathan Sowdhamini

**Author notes:** **Corresponding authors: 1. Email:** (RS), **2. Email:** (GA).

## Abstract

The olfactory system is capable of detecting and distinguishing thousands of environmental odorants that play a key role in reproduction, social behaviours including pheromones influenced classical events. Membrane secretary odorant binding proteins (OBPs) are soluble lipocalins, localized in the nasal membrane of mammals. They bind and carry odorants within the nasal epithelium to putative olfactory transmembrane receptors (ORs). While the existence of OBPs and their significant functions are very well known in insects and laboratory mammals, there is little information about the species-specific OBPs in buffaloes. In fact, the OBP of nasal epithelium has not yet been exploited to develop a suitable technique to detect estrus which is being reported as a difficult task in buffalo. In the present study, using molecular biology and protein engineering approaches, we have cloned six novel OBP isoforms from buffalo nasal epithelium (bnOBPs). Furthermore, 3D model was developed and molecular-docking, dynamics experiments were performed by *In-silico* approach. In particular, we found four residues (Phe104, Phe134, Phe69 and Asn118) from OBP1a, which had strong binding affinities towards two sex pheromones, specifically oleic acid and p-cresol. We expressed this protein in *Escherichia coli* to examine its involvement in the sex pheromone perception from female buffalo urine and validated through fluorescence quenching studies. Interestingly, fluorescence binding experiments also showed similar strong binding affinities of OBP1a to oleic acid and p-cresol. By using structural data, the binding specificity is also verified by site-directed mutagenesis of the four residues followed by in-vitro binding assays. Our results enable to better understand the functions of different nasal epithelium OBPs in buffaloes. They also lead to improved understanding of the interaction between olfactory proteins and odorants to develop highly selective biosensing devices for non-invasive detection of estrus in buffaloes.

## Introduction

Odorant-binding proteins (OBPs), a class of soluble proteins abundantly secreted into the nasal mucus of vertebrates and in the lymph of chemo sensilla in insects, involved in first line of olfaction as carriers of hydrophobic odorants and pheromones [1–4]. The vertebrate OBPs being as a single class of soluble polypeptides, comprises four structurally different families, while two have been identified in the insects named as OBPs and CSPs (chemosensory proteins). Despite the common term, vertebrate OBPs being structurally distinguish with those insects in the terms presenting the typical β-barrel folding in vertebrates and insect OBPs build instead of α-helical segments. [5–10]. Recent studies revealed that Niemann-Pick type C2 proteins (NPC2) from arthropod bears close evolutionary relationship with insect OBPs and CSPs [10, 11]. In vertebrates, odorant binding proteins are belonging to super extracellular proteins called lipocalins [12], which is a key transporter for delivering retinol and fatty acids in the entire body for organism development and differentiation [13, 14]. The vertebrate OBPs share structural similarity with lipocalins. Its compact structure have maximum sequence identity with highly conserved structure of ®-barrel, a sort of cup made of eight antiparallel ®-sheets open one side with 〈-helices at both ends enclosing an internal hydrophobic ligand-binding site [15–18, 3, 19–22].

Most of the studies suggested that odorant binding proteins are pheromone carriers, either in the certain biological gland or secretary body fluids like urine, saliva, seminal fluid etc. [18,23–27] as well as also act as a perceiver of olfactory epithelium [17, 24, 28, 21]. Indeed, it has been well established that, these proteins respond to airborne stimuli of pheromones in olfactory systems and trigger adaptive behavioural responses and/or elicit physiological processes [29–31] and clearly suggesting a common function to deliver the chemical messengers from the site of production to the external environment. In the biological system, the function of lipocalins like vertebrate OBPs is in the interface between external environments and membrane olfactory receptors. The specific biophysical studies revealed that OBPs are considerably stable to pH range from 4.0 to 7.5 [32, 33] and more stable against proteolysis and as well as solvent denaturation compared to odorant receptors. Moreover, presence of ligands inside the binding pocket of OBP further increases its thermal stability and seems like a promising attractive feature for designing suitable sensor for biotechnological applications including variety of transducers, gas sensor, quartz crystal microbalance (QCM), surface acoustic wave devices, organic field effect transistors [34–38]. It has also been proposed that, OBPs are expressed in both the nasal mucus and saliva to play important roles in odor perception and sexual communication in buffaloes [26, 28]. However, the detailed interaction mechanism between pheromones and OBPs, binding site information as well as structural changes induced by pheromone binding has still remained unclear.

Buffaloes, being livelihood of farmers, they contribute much to economy by providing milk, meat, hides and also through agriculture [39, 40]. Yet, the detection of estrus signs, called ‘silent heat’ in buffaloes, is a challenging task which affects their breeding capacity. Silent heats, coupled with poor visual signs of estrus, are the key factors which obstruct the reproductive performance in buffaloes [41]. It is estimated that 50% failure to detect estrus is unpredicted. In the present scenario, different estrus detection methods were established with limited success rates [42, 43]. These current protocols to find estrus are not always reliable, labor-intensive, and need skill and experience. Under natural situation, using the olfactory sense buffalo bull is capable to detect the estrus phase accurately and matting proceeds, in that case the conception rate is 100%. It simply shows that the pheromone molecules released from female estrus are rightly identified by bull. Nevertheless, pheromone based kit developed by us is effective to detect estrus in buffalo about 60% success [44], However, odorant proteins are very ideal elements and probably best choice as sensing elements to detect the pheromones, which are most promising agent in estrus indicator. Therefore, it is aimed to develop a portable, sensor for detecting estrus specific pheromones under field conditions, which would lead to detect the accurate time of animal ovulation phase so that the farmers can take precise decisions to improve the animal breeding.

Based on the reports in buffalo estrus, our long term objective is to construct a biosensor to detect estrus accurately for the effective artificial insemination in buffaloes. The OBP identified and verified in the nasal mucus of buffalo [28] is considered to be the right choice to use this as template to develop biosensor for buffalo estrus detection. Hence, the ultimate aim of this study, to understand of olfaction mechanisms by molecular, biochemical, structural and functional approaches used to determine whether these odorant binding proteins are assisted in perception of sex specific pheromones during the estrus. In order to construct the sensor, the computational and biophysical studies are critically required to know the structural insights and function properties of OBPs. Moreover, 3D model was developed and molecular-docking, dynamics experiments were performed by *In-silico* approach. Furthermore, we predicted structures of bnOBP isoforms and examine their molecular interactions with the two sex pheromones by biophysical methods. Interestingly, fluorescence binding experiments indicated that OBP1a had strong binding affinities towards two sex pheromones specifically oleic acid and p-cresol. Moreover, the key ligand binding residues were confirmed using site-directed mutagenesis.

Here, we aim to (i) identify the putative odorant binding protein from nasal epithelium of buffaloes by cloning, (ii) to understand the structure and function of the nasal epithelium OBPs by biochemical and *in-silico* approach and, (iii) to investigate their molecular interactions and binding efficiency with buffalo estrus-specific pheromones by fluorescence quenching assays and mutational studies.

## Results

### bnOBPs cloning and sequence analysis

The genome of the buffalo recently has been sequenced [45, 46], but its annotation is still not complete. Here, we obtained six different full-length genes from nasal epithelium described in materials and method section. Topo OBP cloning yielded six different protein products with star and stop codons and the ProtParam tool employed to analyse the amino acid sequence and physico-chemical properties **(Table 1).** The sequence comparison of the six bnOBPs with counterparts from other mammals showed high similarities, thus suggesting prime function in pheromone perception. However, Blastp search and phylogenetic tree analyses indicated that this protein had 92% amino acid sequence identity with a bovine OBP protein from *Bos Taurus* (PDB No. 1OBP_A) and belonged to odorant binding proteins family which is well reported [15]. Altogether, their significant similarities with OBPs from other species in National Centre for Biotechnology Information (NCBI) database, the six deduced protein sequences will be deposited in the GenBank database. The six different lengths of open reading frame (ORF) genes encodes that the mature bnOBPs that was expressed by using a pET28a+ vector fused with a His-tag fragment in high yields with soluble forms. The SDS-PAGE analysis showed the highly purified protein as a single band with the molecular weight of approximately around 20 KDa.

**Table 1:** Physicochemical Properties and Amino acid Analysis.

| S. No | Protein ID | Sequence length | Molecular weight (KDa) | Isoelectric point | The grand average of hydropathicity (GRAVY) |
| --- | --- | --- | --- | --- | --- |
| 1. | OBP1a | 174 | 20 | 5.89 | -0.644 |
| 2. | OBP1b | 162 | 19.5 | 5.78 | -0.673 |
| 3. | OBP1c | 174 | 20 | 5.89 | -0.648 |
| 4. | OBP1d | 168 | 19.3 | 5.89 | -0.731 |
| 5. | OBP1e | 174 | 20 | 5.71 | -0.670 |
| 6. | OBP1f | 168 | 18.7 | 5.78 | -0.684 |

### Multiple Sequence Analysis (MSA)

#### Residues highlighted in red boxes are Phe69, Phe104, Asn118 and Phe134

Based on the multiple sequence alignment, isoforms were assigned names accordingly (**Figure 1**). Multiple sequence alignment of bnOBP isoforms shows high degree of conservation of residues across isoforms as well as in comparison with bovine OBP, 1OBP (**Supplementary Table 1**). Differences of few residues among isoforms are observed.

**Figure 1:**
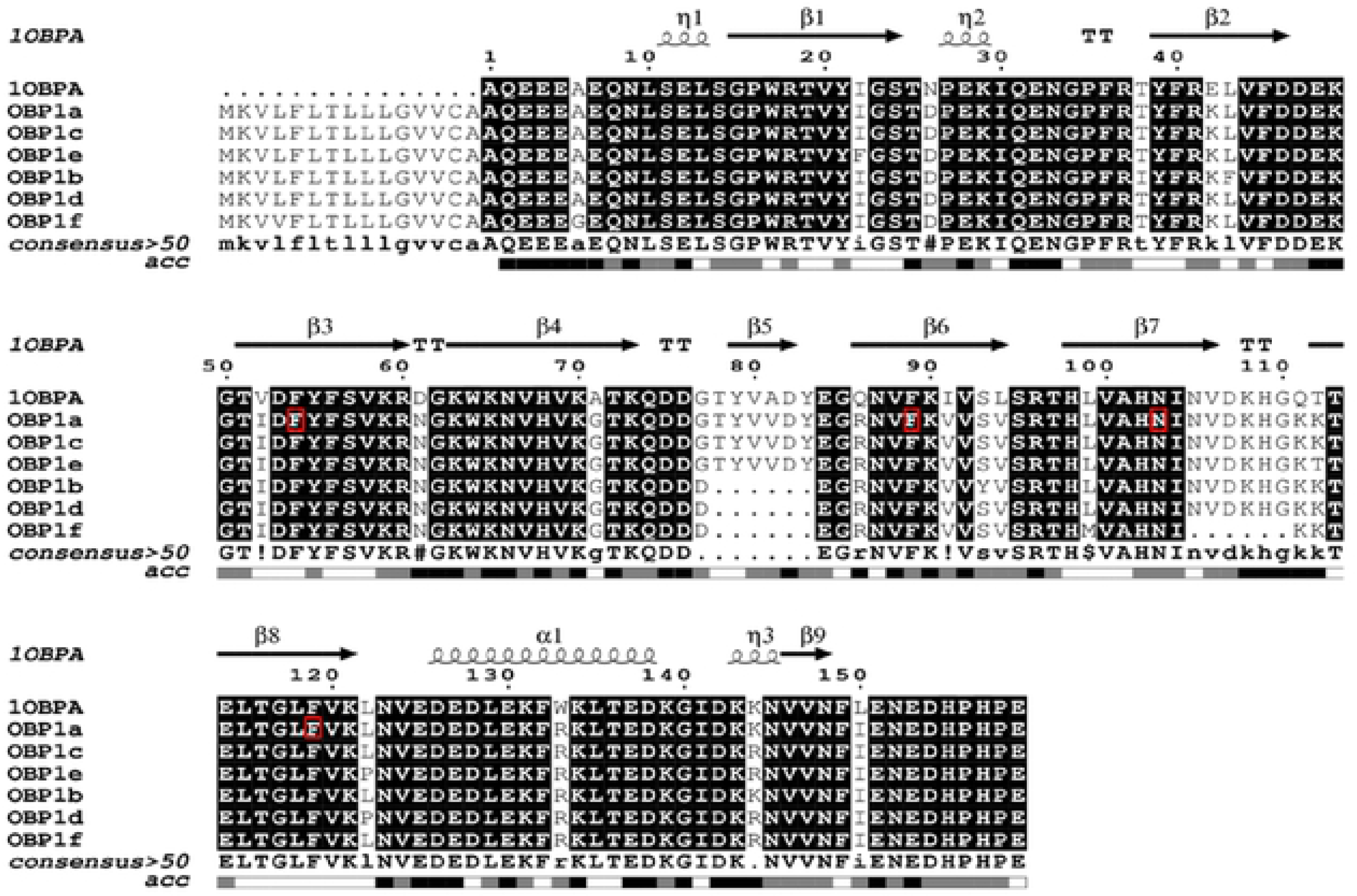
**Multiple sequence alignment of nasal epithelium six OBP isoforms (bnOBPs).** Residues Phe69, Asn 118, Phe 104 and Phe134 corresponding to OBP1a used for site-directed mutagenesis have been marked in red rectangular boxes.

### Domain analysis

All six isoforms were positive for the presence of lipocalin domain from the lipocalin/cytosolic fatty-acid binding protein family (PF00061.23) with high confidence. Lipocalin folds are characteristic of vertebrate OBPs (**Supplementary Table 2**).

### Homology modelling

In the present study, protein nasal mucus segregated protein from *Bos taurus* [PDB: 1OBP] shared the maximum sequence identity with bnOBPs and was further chosen as the template for homology modelling. Additionally, it appeared as co-clusters and has maximum evolutionary relationship with buffalo genome [28]. The constructed models were further validated by Pro-CHECK software and visually inspected by PyMOL software tool (**Figure 2**). The superposition of the models and the template confers that the structures and folding patterns were very highly similar and the root mean square deviation (RMSD) values are listed in **Table 2**. Put together, the predicted models bnOBPs were found to be structurally reasonable and reliable.

**Figure 2:**
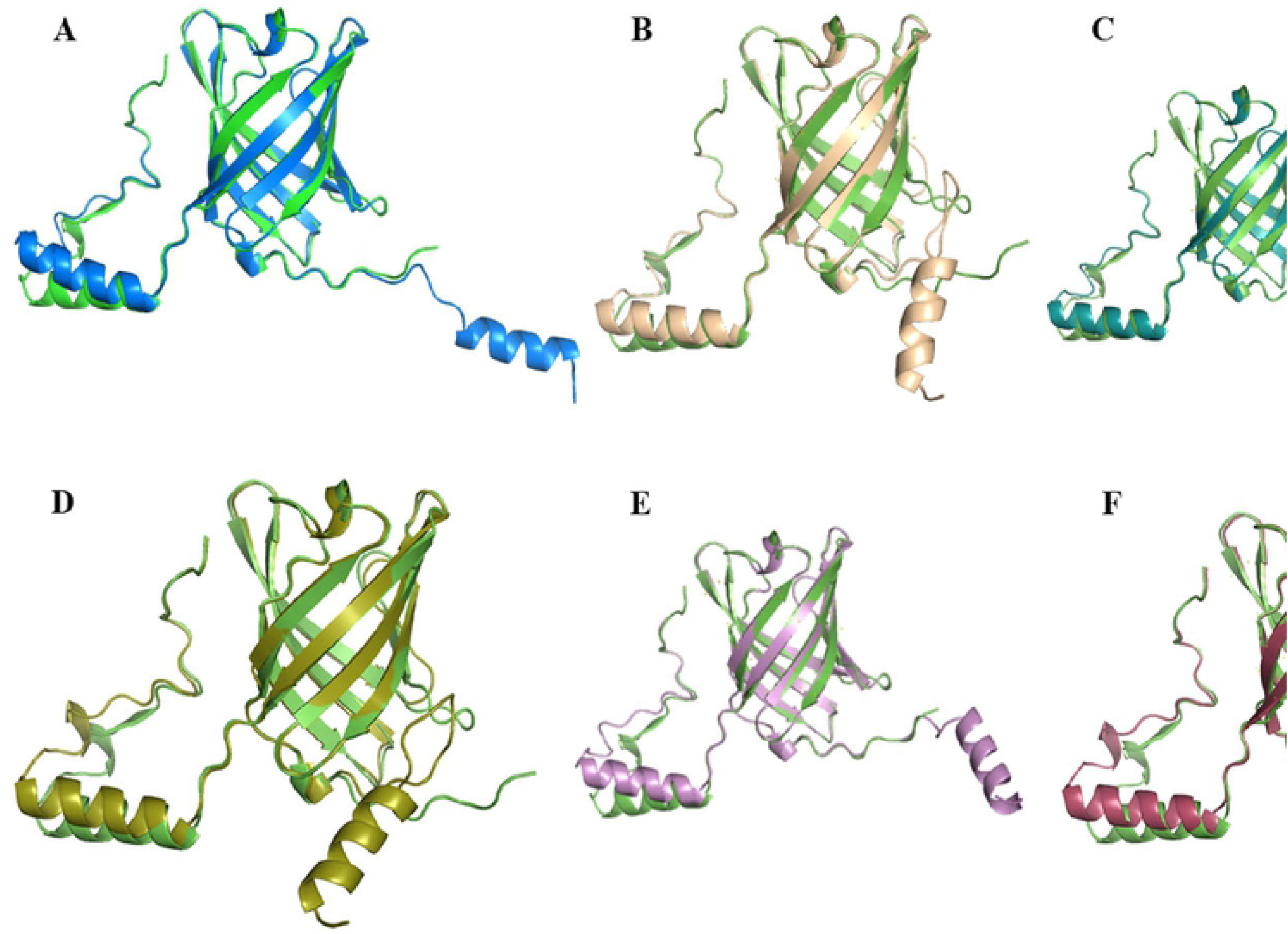
Structural Superimposition of nasal epithelium six OBP isoforms. (A-F) The crystal structure of bovine 1OBP (template) is green colour and all other nasal epithelium OBPs (queries) shown different colours respectively.

**Table 2:** The rationality values and 3D structure verification scores of the constructed models.

| Isoforms | Sequence identity with template PDB (1 OBP) % | Ramachandran Plot analysis |  | Root mean square deviation (RMSD) value |
| --- | --- | --- | --- | --- |
|  |  | Favoured regions % | Residues in additional allowed regions % |  |
| OBP1a | 91.82 | 90.3 | 9.7 | 0.199 Å |
| OBP1b | 86.79 | 88.9 | 9.0 | 0.438 Å |
| OBP1c | 91.14 | 93.5 | 5.8 | 0.974 Å |
| OBP1d | 81.65 | 87.2 | 9.4 | 1.075 Å |
| OBP1e | 87.34 | 90.9 | 8.4 | 1.424 Å |
| OBP1f | 82.39 | 89.3 | 10 | 0.284 Å |

### Site analysis

#### Binding sites on Isoform OBP1a

OBP1a has been isolated from the buffalo nasal epithelium. It has eight anti-parallel beta strands enclosing a beta barrel structure centrally and flanked by two terminal alpha helices connected to the beta barrel by a loops. The C-terminus has a loop and is preceded by one of the alpha-helices. These features at the structural level have been retained by isoforms 1a to 1f. There were 5 potential sites predicted for isomer OBP1a, out of which one site was a strong ligand-binding site (pocket 1) and the second ranked site was a partial binding site (pocket 2) (**Figure 3**). Pocket 1, also a classical ligand-binding site, was located as a central pocket in the hydrophobic cavity of OBP1a (**Figure 3B**). The second site predicted, Pocket 2, is an allosteric site and is located laterally between the beta barrel and C-terminal side of the structure (**Figure 3C**). **Throughout the study the central cavity will be referred to as pocket 1 (or P1), and the lateral ligand-binding site will be referred to as pocket 2 (or P2).** The sites predicted that were shortlisted further based on favourable ligand-binding and druggability criteria (**Table 3**). Docking and affinity results suggested oleic acid binds selectively to isoforms OBP1a, OBP1b and OBP1d at the central cavity whereas p-cresol docks at the central cavity across isoforms OBP1a-f (**Tables 4 and 5**). Residue Phe69 interacts with p-cresol through pi-pi stacking interactions, whereas Asn118 forms a hydrogen bond with the ligand (**Figure 4**) when docked in pocket 1.

**Figure 3:**
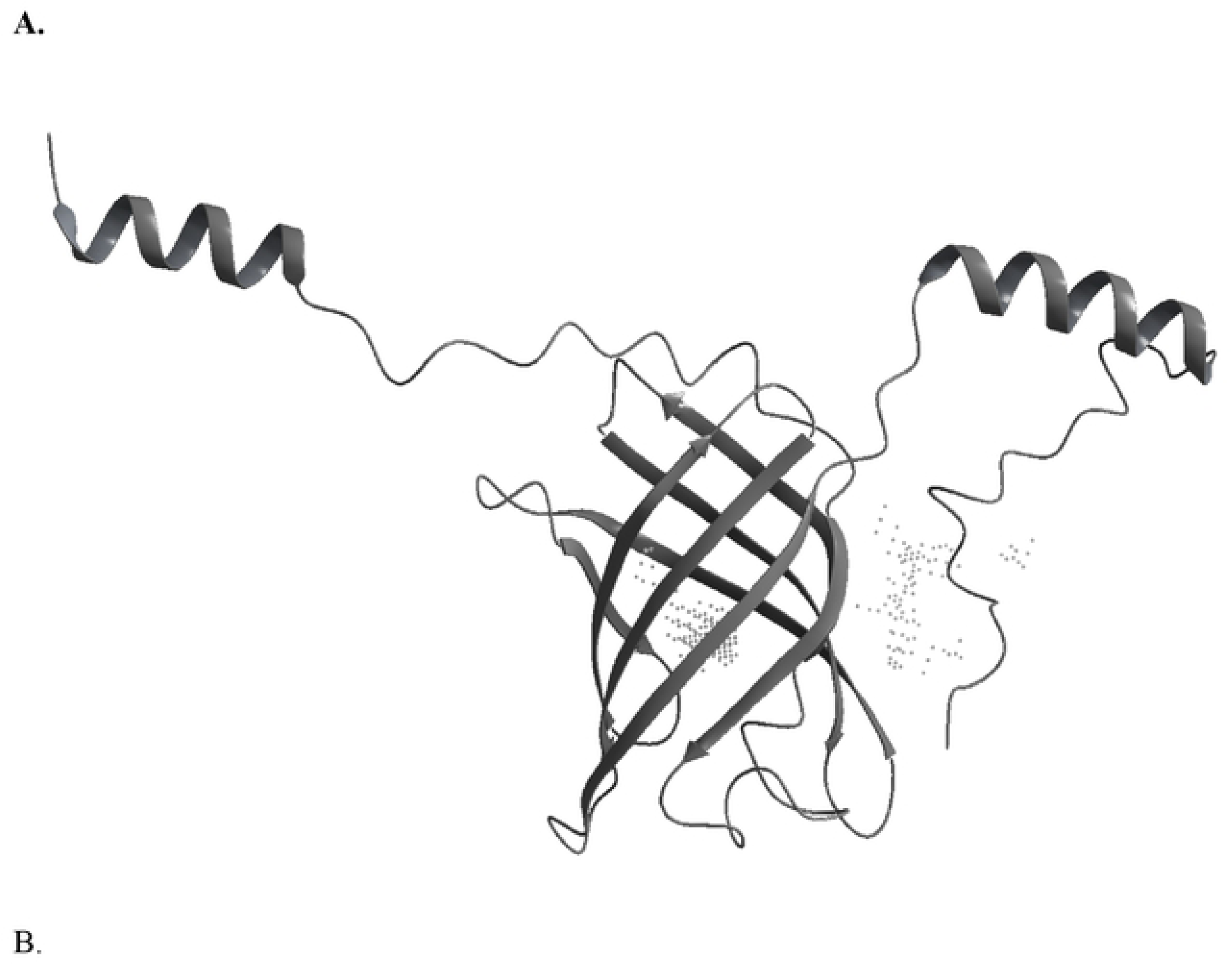
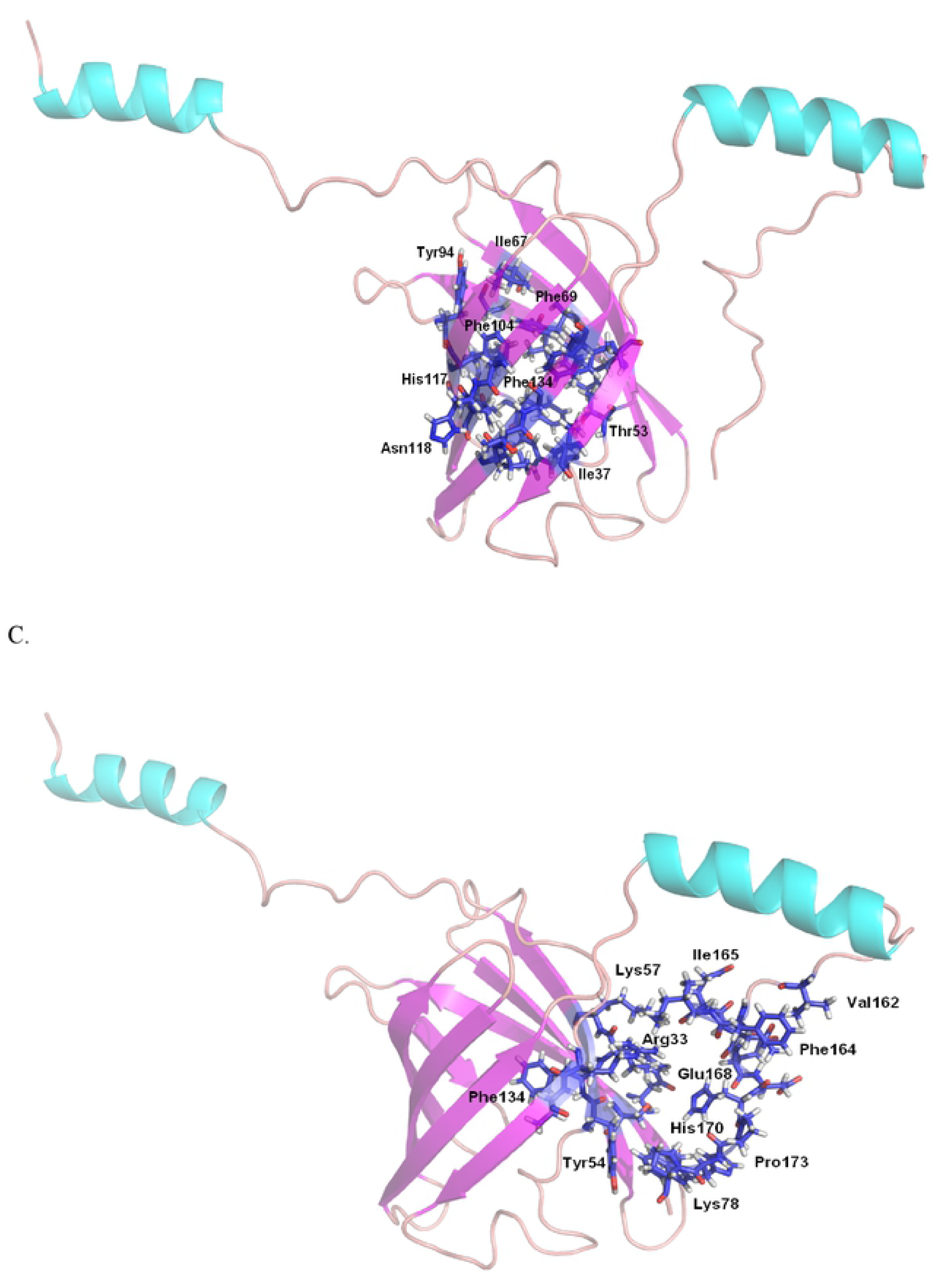
Druggable sites on OBP1a isoform A. Pockets depicted as dotted sheres, B. Residues of Pocket 1 (central cavity) depicted as sticks in dark blue and C. Residues of Pocket 2 (laterally located) depicted as sticks in dark blue.

**Figure 4:**
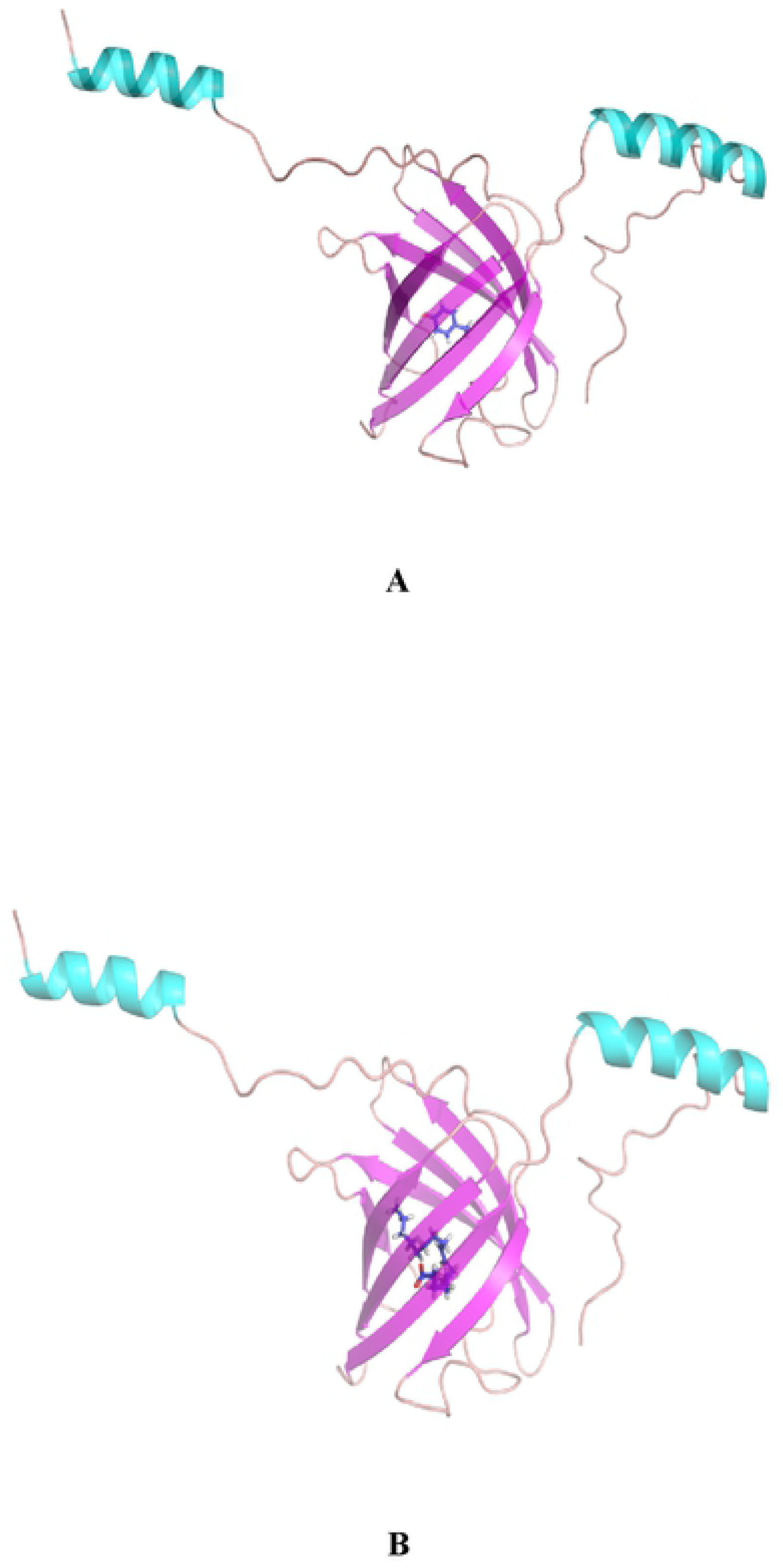
**Protein-ligand docking of OBP1a with (A) p-cresol and (B) oleic acid at the central binding cavity.** Residue Phe69 interacts with p-cresol through pi-pi stacking interactions, whereas Asn 118 forms a hydrogen bond with the ligand (Figure 4) when docked in pocket 1.

**Table 3:** Potential ligand-binding sites in buffalo OBP isoforms Protein-ligand docking and affinity for wild type isoforms-ligand complexes

| Isoform | Cavity location | Site ID | Ligand-binding score | Druggability | Size (Å<sup>2</sup>) | Volume (Å<sup>3</sup>) |
| --- | --- | --- | --- | --- | --- | --- |
| **OBP1a** | **Central** | **1** | **1.35** | **1.46** | **113** | **144.746** |
| OBP1a | Lateral | 2 | 0.92 | 0.94 | 93 | 254.506 |
| **OBP1a F69A** | **Central** | **1** | **1.334** | **1.440** | **134** | **200.998** |
| OBP1a F69A | Lateral | 2 | 0.995 | 1.040 | 116 | 325.164 |
| **OBP1a F104A** | **Central** | 1 | 1.344 | 1.458 | 153 | 217.462 |
| OBP1a F104A | Lateral | 2 | **0.972** | **0.945** | **97** | **236.67** |
| **OBP1a N118A** | **Central** | 2 | 1.36 | 1.475 | 110 | 119.364 |
| OBP1a N118A | Lateral | 1 | **0.966** | **1.008** | **153** | **357.749** |
| **OBP1a F134A** | **Central** | 1 | 1.334 | 1.440 | 147 | 197.911 |
| OBP1a F134A | Lateral | 2 | 0.984 | 1.011 | 119 | 270.627 |
| **OBP1f** | **Central** | 3 | **1.257** | **1.312** | **94** | **120.393** |
| OBP1f | Lateral | 2 | 1.103 | 1.007 | 116 | 211.288 |
| OBP1f | Lateral | 1 | 0.982 | 1.031 | 131 | 366.667 |
| **OBP1e** | **Central** | 2 | **1.297** | **1.395** | **84** | **111.818** |
| OBP1e | Lateral | 1 | 0.992 | 1.058 | 109 | 268.569 |
| **OBP1b** | **Central** | 2 | **1.2165** | **1.3079** | **140** | **195.853** |
| OBP1b | Lateral | 1 | 1.016 | 1.033 | 216 | 405.083 |
| OBP1b | Lateral | 3 | 0.987 | 1.025 | 114 | 344.715 |
| **OBP1d** | **Central** | 2 | **1.2318** | **1.2859** | **138** | **223.293** |
| OBP1d | Lateral | 1 | 1.1214 | 0.9707 | 156 | 298.067 |
| OBP1d | Lateral | 3 | 1.002 | 1.043 | 119 | 322.077 |
| **OBP1c** | **Central** | 2 | **1.374** | **1.500** | **112** | **131.369** |
| OBP1c | Lateral | 1 | 0.9808 | 1.019 | 130 | 318.990 |
| OBP1c | Lateral | 3 | 0.816 | 0.8006 | 71 | 180.761 |

**Table 4:**
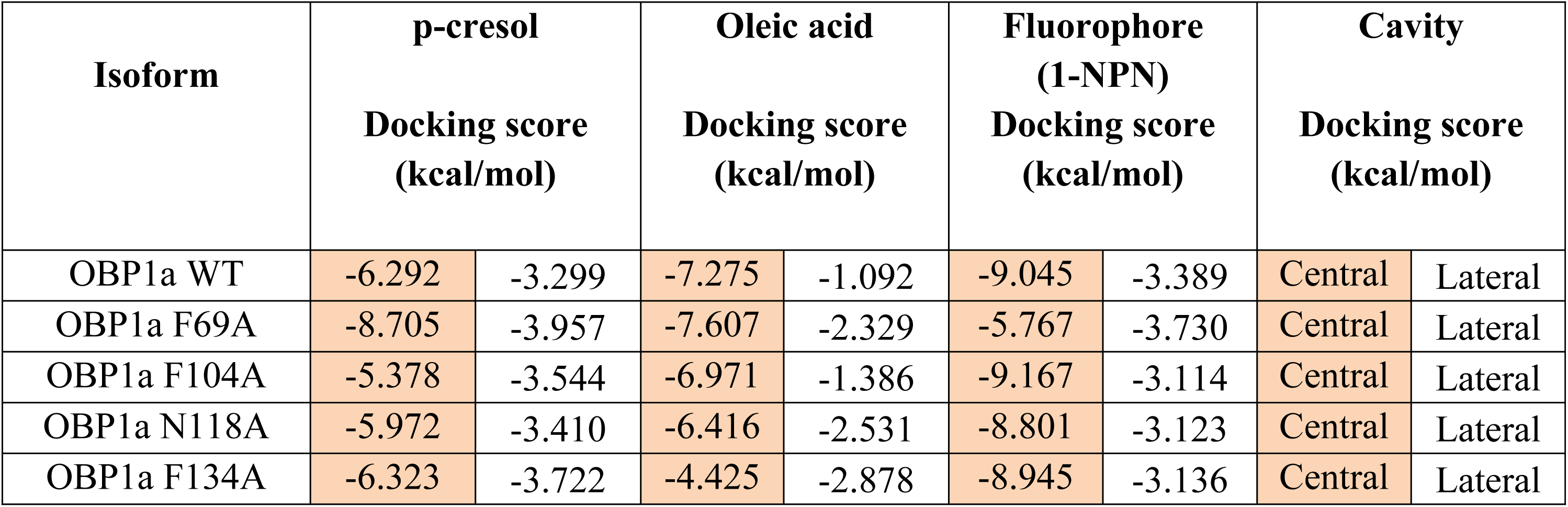
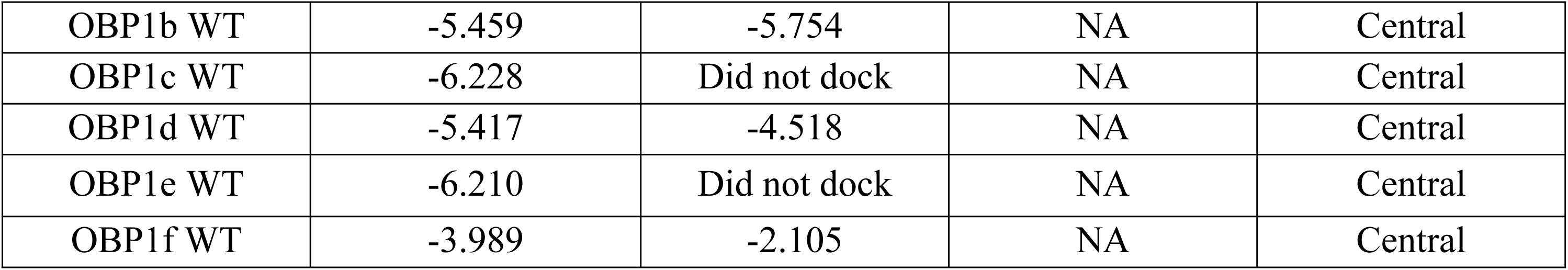
Scores for top poses of protein-ligand docked structures

**Table 5:** Affinity scores of docked protein-ligand complexes

| Isoform | Cavity |  | p-cresol |  | Oleic acid |  | 1-NPN |  |
| --- | --- | --- | --- | --- | --- | --- | --- | --- |
| **OBP1a** | Central | Lateral | -36 | -26.81 | -9.27 | -29.82 | -50.29 | -38.64 |
| **OBP1a F69A** | Central | Lateral | -35.21 | -32.57 | -29.44 | -36.24 | -51.85 | -38.84 |
| **OBP1a F104A** | Central | Lateral | -33.85 | -34.18 | -13.11 | -33.79 | -54.22 | -35.14 |
| **OBP1a N118A** | Central | Lateral | -35.58 | -24.59 | 38.23 | -43.92 | -49.59 | -41.23 |
| **OBP1a F134A** | Central | Lateral | -38.09 | -33.55 | -31.47 | -45.68 | -54.81 | -33.55 |
| **OBP1b** | Central |  | -30.85 |  | -39.45 |  | NA |  |
| **OBP1c** | Central |  | -35.98 |  | No pose |  | NA |  |
| **OBP1d** | Central |  | -30.42 |  | -19.9 |  | NA |  |
| **OBP1e** | Central |  | -35.57 |  | No pose |  | NA |  |
| **OBP1f** | Central |  | -34.1 |  | 5.65 |  | NA |  |

### OBP1a WT

Molecular dynamics and RMS deviation of the OBP1a wild type model suggests a convergence at 50 ns onwards (**Figure 5A**) with residues Glu63, Asn76, Gly77, Asp90-91 showing moderate level of RMS fluctuation (**Figure 5B**) whereas His124 (3.48 Å), Glu142 (4.8 Å), Asp153 (6.73 Å) show steadily increasing levels of fluctuation. Presence of alpha helix near the C-terminus is consistent throughout the course of the simulation although the N-terminal helices remain stable less than 80% of the time course of 100 ns (**Figure 5C**).

**Figure 5:**
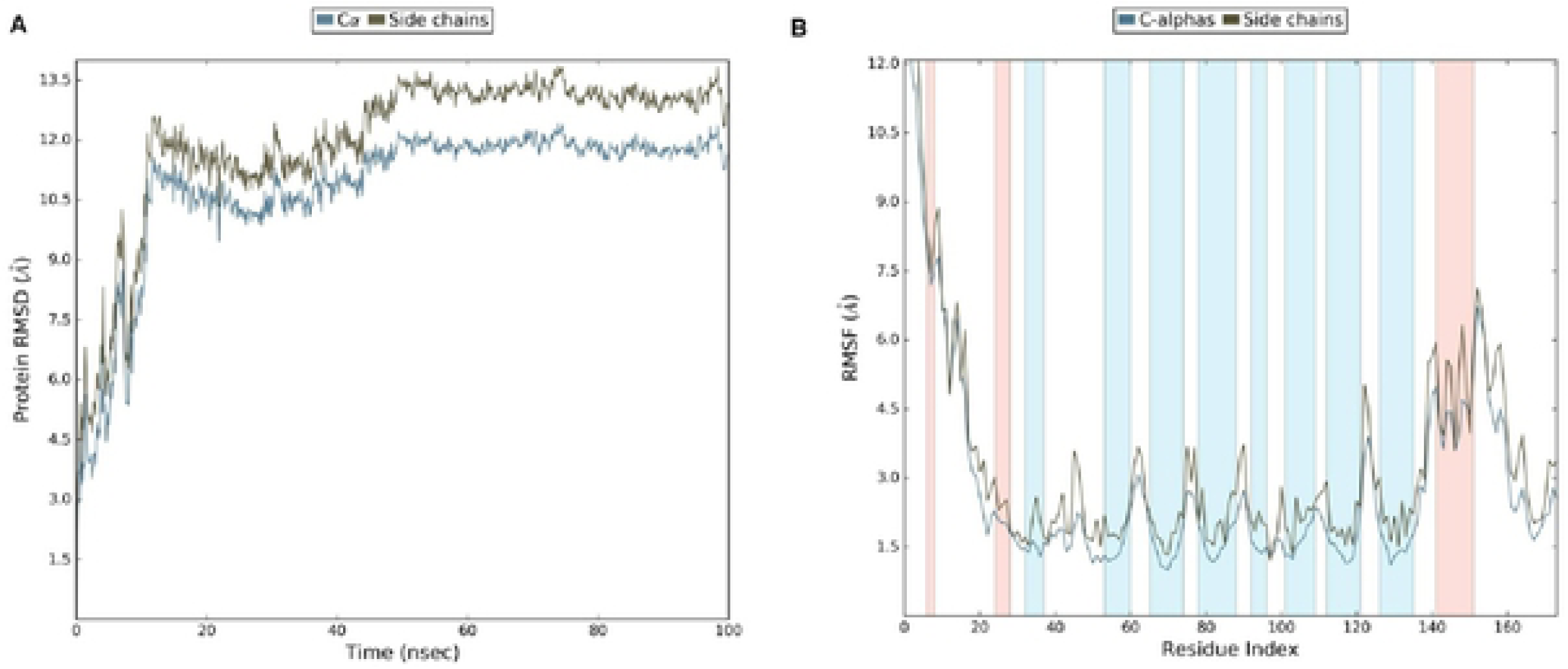
**Molecular dynamic simulation of apo-OBP1a WT for 100 ns** (A) RMSD of the C-alpha atoms of comparatively modelled bovine OBP structure (B) RMSF of the backbone of protein.

**Figure 5:**
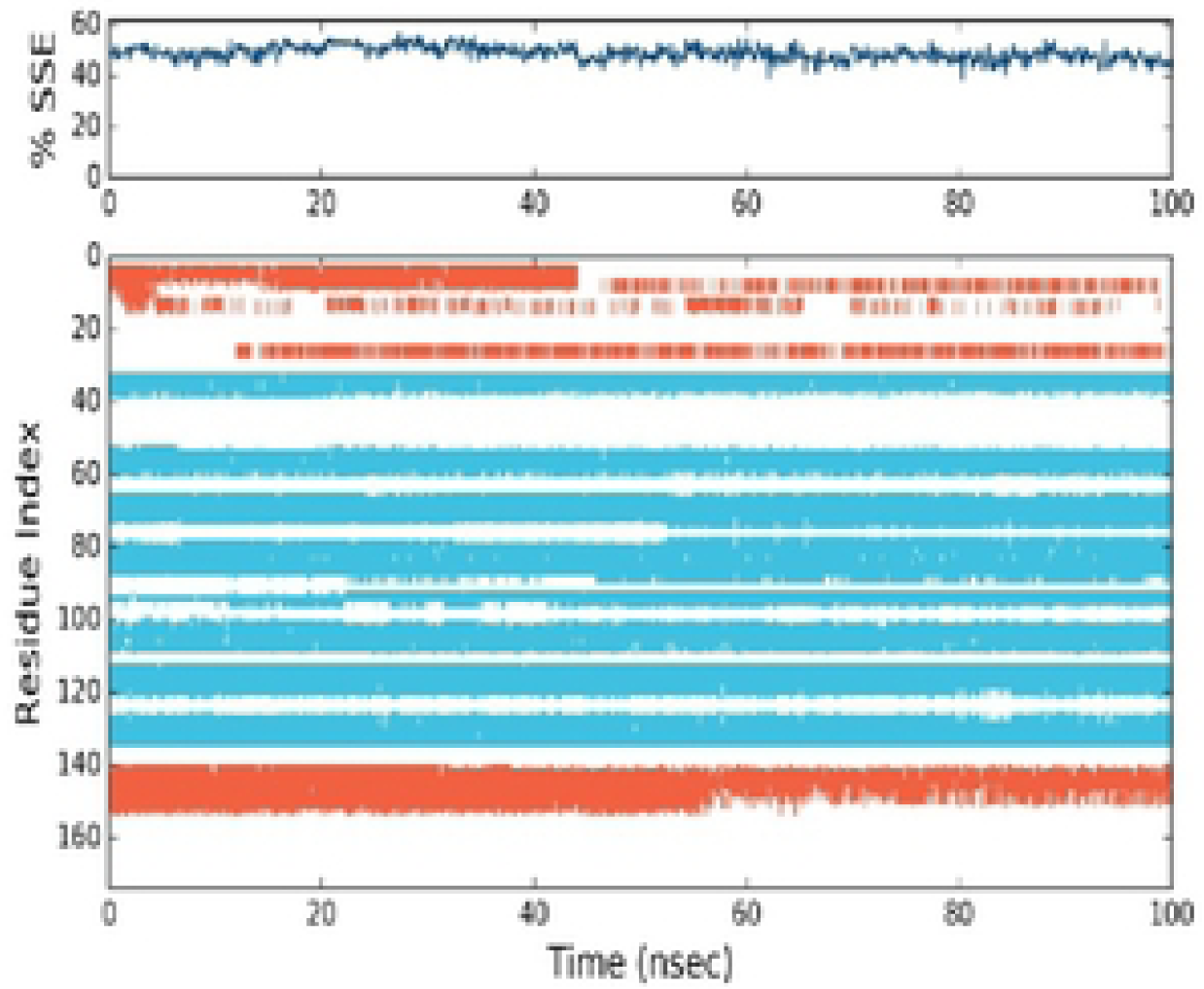
**Molecular dynamic simulation of apo-OBP1a** (C) Secondary structure stability across 100 ns. Presence of alpha helix near the C-terminus is consistent throughout the course of the simulation although the N-terminal helices remain stable less than 80% of the time course of 100 ns **(Figure 5C)**.

### Molecular Dynamic simulation analysis for isoform OBP1a with p-cresol and oleic acid

The **RMSD** of the MD simulation trajectory of OBP1a with p-cresol suggests that the protein-ligand complex attains convergence after **20 ns** from the simulation start-point and remains steady atleast upto 100 ns from the start-point (**Figure 6A**) suggesting that **OBP1a-p-cresol complex at the central binding cavity would reach equilibration conditions faster than the wild type isoform alone (Supplementary Text).**

**Figure 6:**
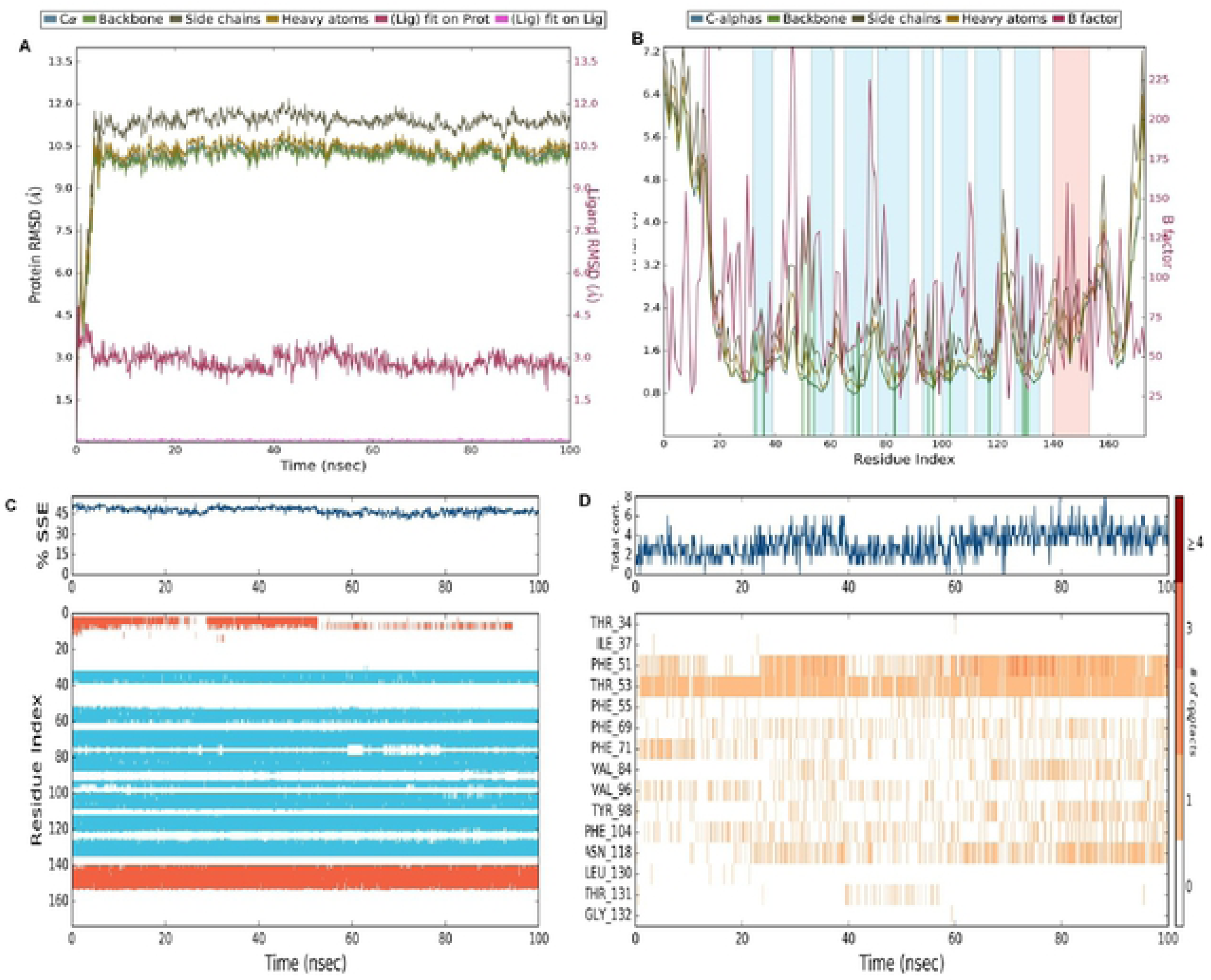
**Molecular dynamic simulation of OBP1a – p-cresol for 100 ns in central cavity** (A) RMSD of the C-alpha atoms of comparatively modeled bnOBP structure and ligand (B) RMSF of the backbone of protein (C) Secondary structure stability of the complex (D) Residue contacts with ligand.

The RMSF of the residues suggests that the C-terminus tail that is a loop shows highly increased fluctuations from residue Glu166 to Glu174 (**Figure 6B**). This is similar to that seen in the first 20 residues suggesting **that alpha helices near both termini show high levels of fluctuation in the presence and absence of p-cresol.**

The first 10 residues are part of an alpha helix about 40% of the time whereas the helix is observed for residues 141 to 154 throughout the simulation. Eight **beta strands** prevail throughout the duration of the simulation for sets of residues 33-40, 54-62, 66-76, 78-89, 94-98 101-110, 113-122 and 127-136 while the remaining residues are in flexible loop regions (**Figure 6C**). Thr53 interacts with the ligand directly via hydrogen bond upto ∼31% of the time compared to Phe51, a polar residue, which interacts with p-cresol only ∼2% of the time. However, formation of a **water-bridge** between Thr53 and p-cresol mediates contact for atleast 85% of the simulation, and in case of Phe51 extends contact upto 60% of the time, suggesting a water bridge-mediated mechanism of OBP1a-p-cresol binding at the central cavity in the beta barrel. **Asn118** interacts with ligand not more than 30% strength mostly towards the last 20 ns of the simulation with water bridges accounting for upto 11% of the interactions between 40 to 80 ns (**Figure 6D**). In contrast, Thr53 interacts directly with the ligand upto the first 20 ns and between 40 to 60 ns. Time intervals of 20 - 40 ns and 60 – 100 ns are water bridge-mediated interactions with p-cresol.

**Hydrophobic interactions** were observed with the ligand by residues Phe51, Phe55, **Phe69**, Phe71, Val84, Val96, Tyr98, **Phe104** and Leu130. For residues Phe51, Phe55, **Phe69**, Phe71, Tyr98 and **Phe104**, pi-pi stacking interactions were also observed. However, hydrophobic interactions, overall, did not account for more than 20% interaction strength with the ligand in case of each residue and were sparsely present throughout the time course. There are no ionic interactions indicated during the time course of 100 ns of protein-ligand interactions.

The RMSD of the MD simulation trajectory of OBP1a with oleic acid suggests that the protein-ligand complex starts converging from 20 ns from the simulation start-point and attains convergence between 70ns to 100ns from the start-point (**Figure 7A**).

**Figure 7:**
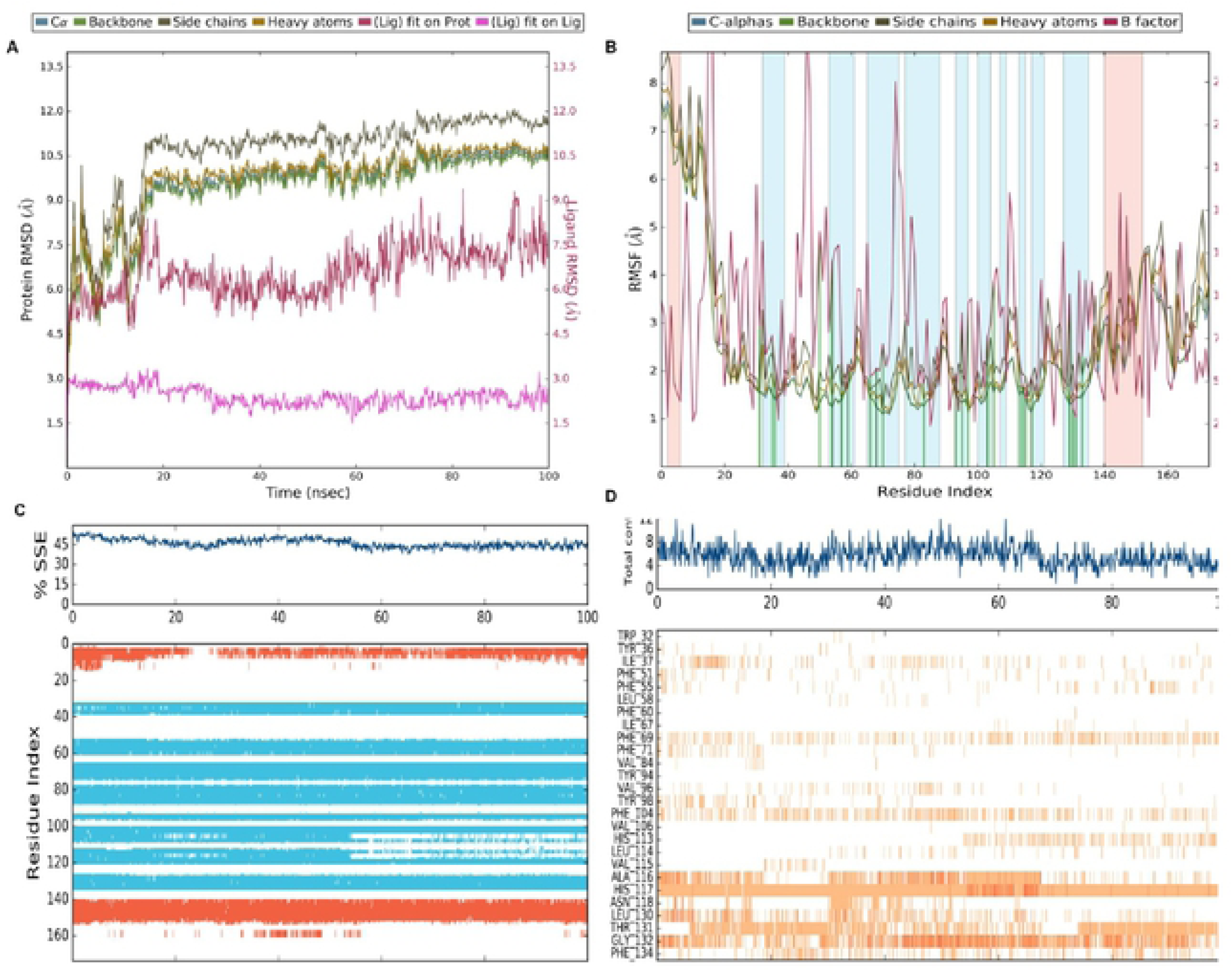
**Molecular dynamic simulation of OBP1a – oleic acid for 100 ns in central cavity (pocket 1)** (A) RMSD of the C-alpha atoms of comparatively modeled bnOBP structure and ligand (B) RMSF of the backbone of protein (C) Secondary structure stability of the complex (D) Residue contacts with ligand.

**Figure 8:**
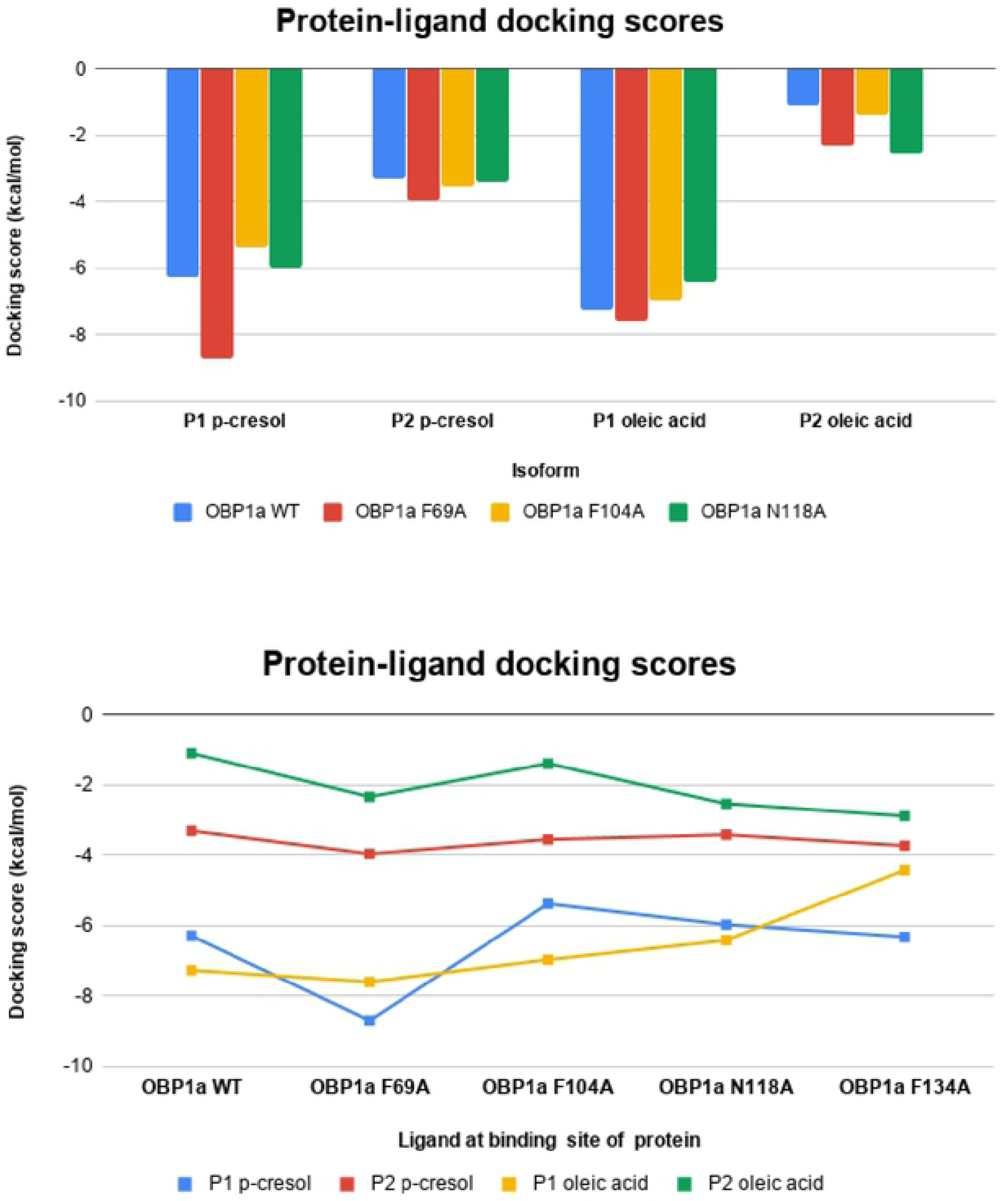
Protein-ligand docking scores across (A) isoforms and (B) binding pockets

**Figure 9:**
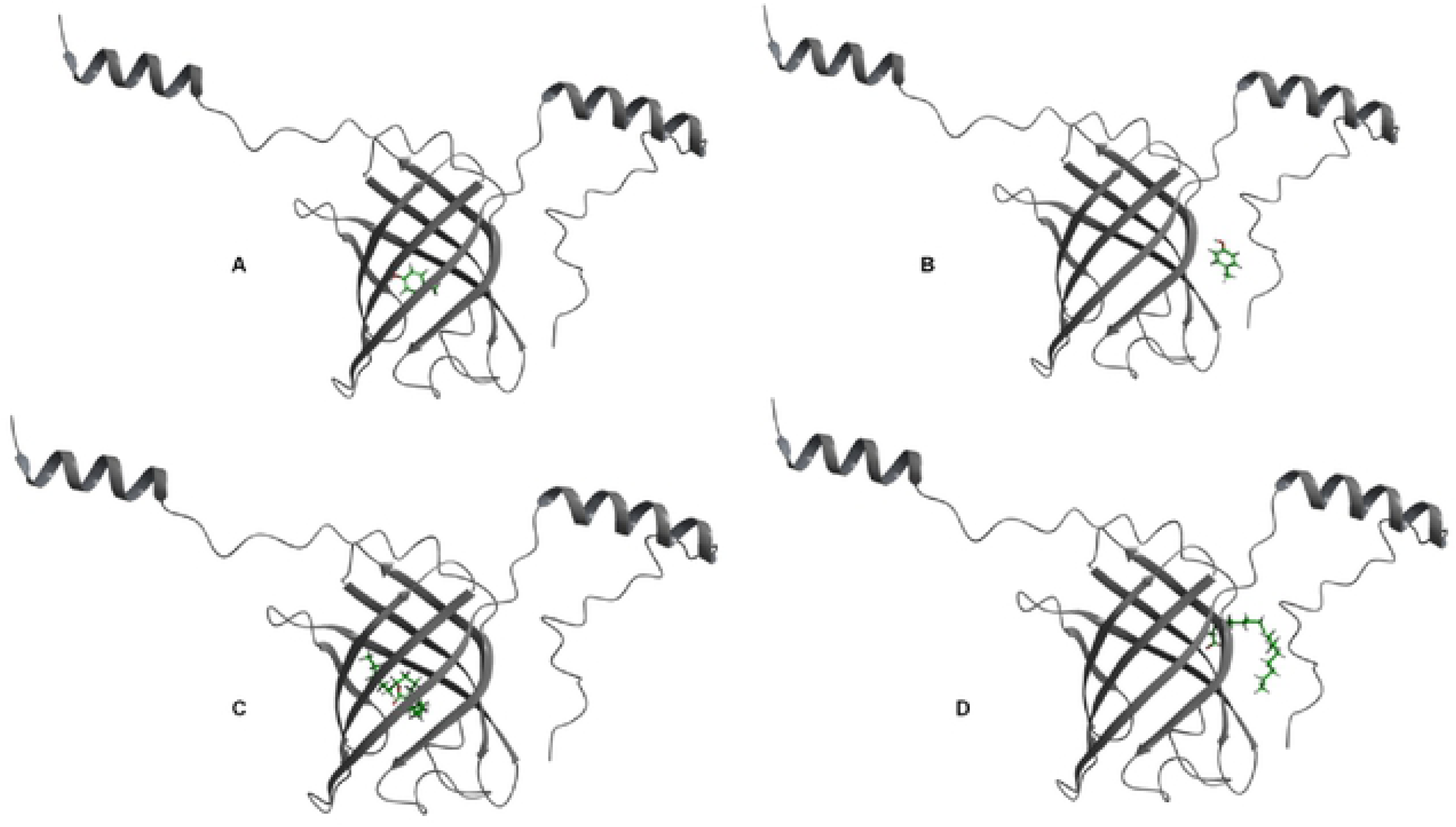
Docked poses for OBP1a isoform with ligands p-cresol (A & B) and oleic acid (C & D) at pocket 1(A & C) and pocket 2 (B & D)

The RMSF of the residues suggests that the C-terminus tail that is a loop shows fluctuation from residue Glu166 to Glu174 which does not exceed 3.8 Å (**Figure 7B**).

Moderate levels of RMS fluctuation are observed in Glu63, Asn76, Gly77, Asp90, Asp91, Arg111, Thr112, Lys123, Asp141, Glu142 and Glu145. However, residues Asp153, Lys154 and Gly155 exceed 4 Å RMSF (4.45 Å). Residues typically exhibited fluctuation of greater than 3 Å for those beyond the first 120 residues.

The first five residues, Val3-Thr7, are part of an alpha helix about 80% of the time whereas helix is observed for residues Asp141 to Lys153 throughout the simulation. **More than eight beta strands** prevail throughout the duration of the simulation for sets of residues with Arg33-Thr40, Tyr54-Asp62, Thr66-Asn76, Lys78-Gln89, Tyr94-Tyr98, Arg101-Lys105, and Asn118-Asp122 prevailing as a beta strand **throughout the course** suggesting changes in the beta barrel structure of the isoform when oleic acid binds at the central pocket. In contrast, Ser108-Ser110 and Leu114-Ala116 are beta strands about 80% of the time (**Figure 7C**).

**Hydrogen bond** between ligand and His117 prevailed throughout and also with residues Gly132 (54.6%), Thr131 (44%), **Asn118** (13%), Ala116 (16%) and His113 (2.6%).

**Water bridges** stabilize Gly132 (57.6%), Thr131 (14.4%), Ala116 (17.7%), His113 (18.2%), His117 (13%) and Leu130 (13.7%). Ala116 water-bridge stabilizes interaction from 30ns to ∼55ns after which H bond between 55 to ∼68ns helps interact further. Gly132 interacts with ligand via water-bridge, as well as hydrogen bonds, throughout the time course but discontinuously. In case of His117, both hydrogen bonding and water-bridge persisted from 55 to 70 ns whereas the water-bridge between 30 to ∼70ns for Thr131 bridging H bond with ligand (**Figure 7D**).

**Hydrophobic interactions** do not involve pi-pi or pi-cation interactions and were observed with **Phe104 (35%), Phe69(30.6%), Phe134 (28.3%),** leu130 (15.5%), Ala116 (10.9%), Ile37(10%), Phe51(3.9%), Phe55(6.4%), Leu58(1.1%), Ile67(1.4%), Phe71(1.6%), Val96(6.9%), Tyr98(2.4%), Val106(0.7%), Leu114(6.6%), Val115(3%), There are no ionic interactions indicated during the time course of 100 ns of protein-ligand interactions.

### Selection of residues for site-directed mutagenesis

Residues that were observed to interact with p-cresol and oleic acid during preliminary MD analysis (**Figures 6, 7, S1 and S2**) were analysed in conjunction with docked poses and their thermal affinity values. The duration and nature of interactions including hydrogen bonds, ionic, water bridges and hydrophobic was taken into account while selecting residues F69, N118, F104 and F134.

Isoform OBP1a was thus mutated to alanine on each of the four residues respectively. Accordingly, four mutant isoforms were constructed **OBP1a F69A, OBP1a N118A, OBP1a F104A and OBP1a F134A**.

### Docking and binding affinity of mutant protein-ligand complexes

Protein-ligand docking yielded complex poses for isoforms of OBP1a and ligands p-cresol and oleic acid respectively. Interestingly, OBP1c and OBP1e did not bind to oleic acid suggesting isoform-ligand binding preference based on geometry and physio-chemical properties (**Figure 7A**).

Among all wild type isoforms, OBP1a showed slightly higher binding affinity to p-cresol (−36 kcal/mol) at pocket 1. OBP1b showed highest binding affinity for oleic acid at pocket 1 (−39 kcal/mol) and OBP1c, 1e and 1f did not bind to oleic acid at pocket 1.

It can be observed that docking scores across OBP1a wild type/mutant isoforms with ligands are significantly lower at pocket 2 than at pocket 1 (**Figure 7B**). This could be explained by the fact that the scoring function used identifies pocket 1 as having higher hydrophobicity, druggability and ligand-binding tendency in general. Across binding pockets, it can be observed that isoforms do not differ significantly in their docking score for a given pocket-ligand pair. There are some exceptions to this trend. OBP1a F69A (−8.71 kcal/mol) has a significantly higher docking score than other complexes at pocket 1 with p-cresol as ligand (**Table 4**). Similarly, OBP1a F134A-oleic acid complex has a significantly low docking score (−4.43 kcal/mol) than other complexes at pocket 1.

Interestingly, mutant forms OBP1a F69A, F104A, N118A and F134A showed similar binding affinities in the range of −33 to −38 kcal/mol to p-cresol at pocket 1 as the wild type suggesting that pocket 1 is highly favored as a binding site for a small aromatic ligand like p-cresol (**Figure 10**). It could be also possible that there would be other strong interactions that arise in a solvent system over a time period that could compensate for the effects due to mutation at the sites F69, F104, N118 and F134 in a centrally enclosed binding location. OBP1a F134A also has slightly higher affinity (−38.09 kcal/mol) for p-cresol at pocket 1 than OBP1a wild type (−36 kcal/mol) and a similar trend is seen for that in pocket 2 as well (**Table 4**).

**Figure 10:**
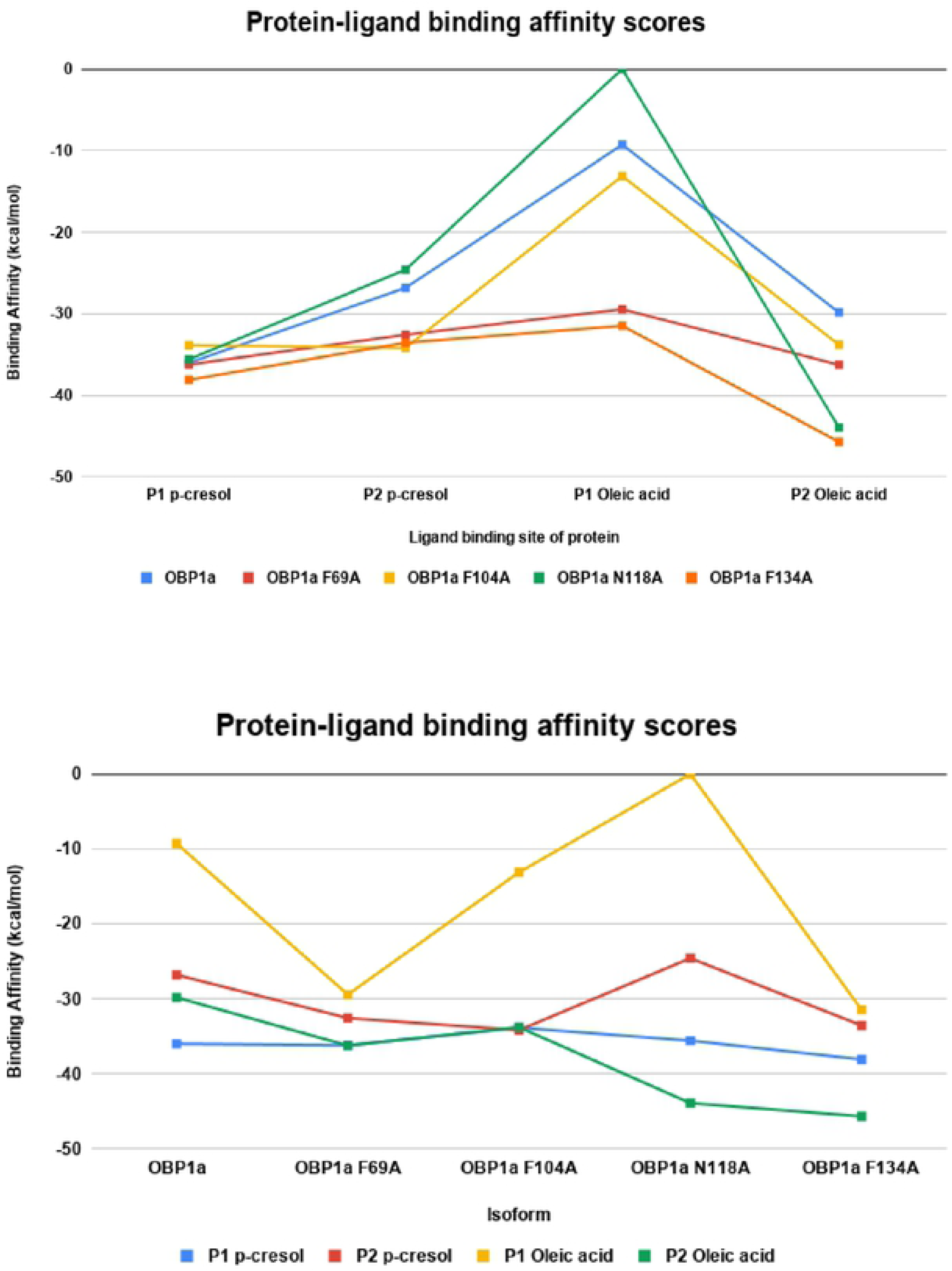
Thermodynamic binding affinity calculated A & B between <u>b</u>nOBPs isoform-ligand pairs at the central binding cavity

### Protein-ligand interactions visualization aka summary of docking

Protein-ligand interactions were analyzed from shortlisted docked poses of protein and ligand. Mutants of OBP1a (F69A and N118A) lose pi-pi stacking interaction with p-cresol but form a hydrogen bond between Thr53 and p-cresol (**Figures 11C and 11D**). Residues Asn118 interacts with oleic acid through hydrogen bond (**Figure 12A**) when docked in pocket 1. Mutant of OBP1a (F104A) loses the hydrogen bond interactions with oleic acid but mutant 134A retains the same hydrogen bond (**Figures S5C and S5D**). However, in case of oleic acid docked at pocket 2, both Arg54 and Asn81 form hydrogen bonds with oxygen atoms of oleic acid (**Figure 12B**).

**Figure 11:**
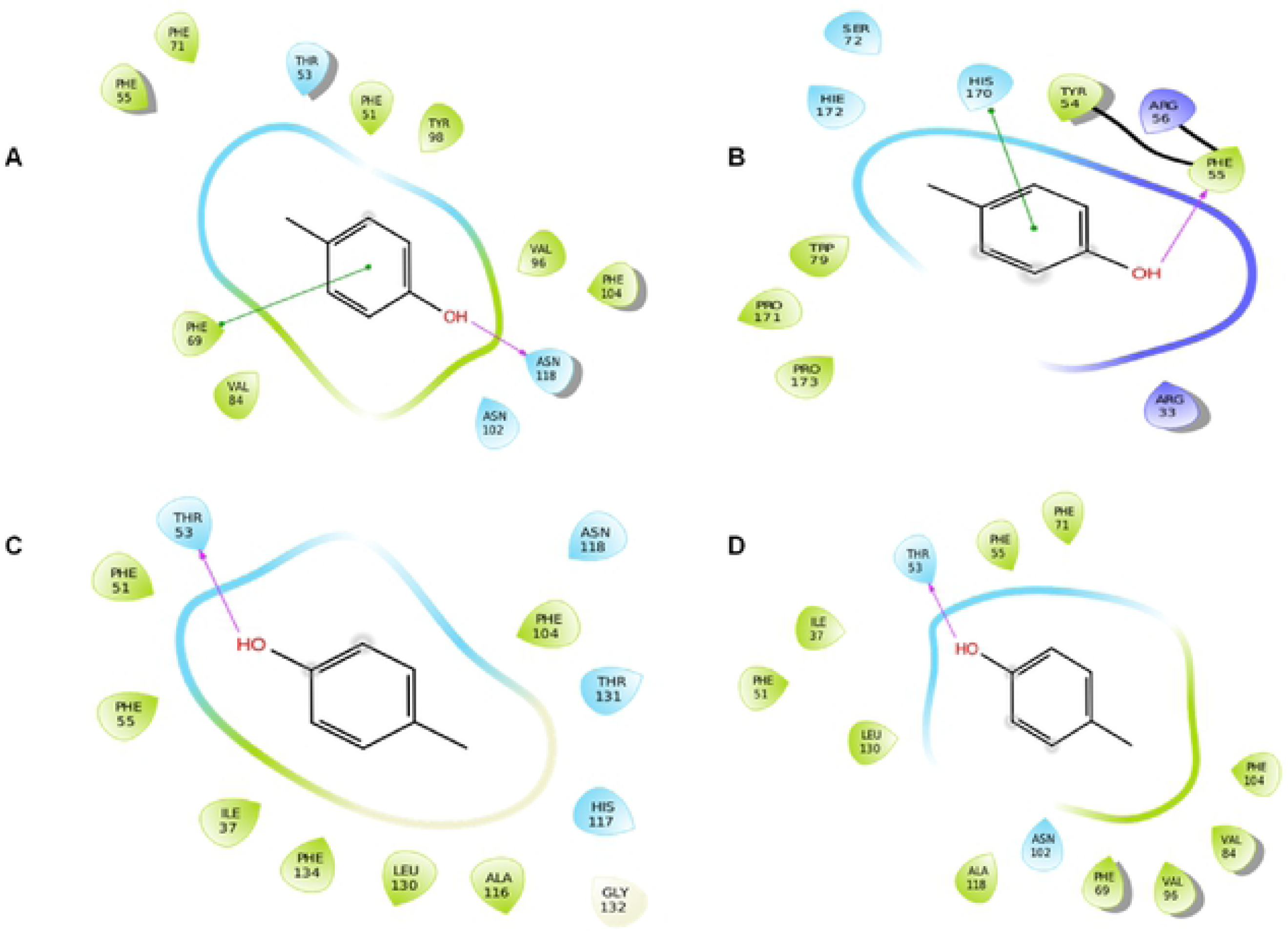
Protein-ligand interactions with p-cresol as ligand docked with OBP1a (A) in pocket 1, OBP1a (B) in pocket 2, OBP1a F69A in pocket 1 (C) and OBP1a N118A in pocket 1 (D).

**Figure 12:**
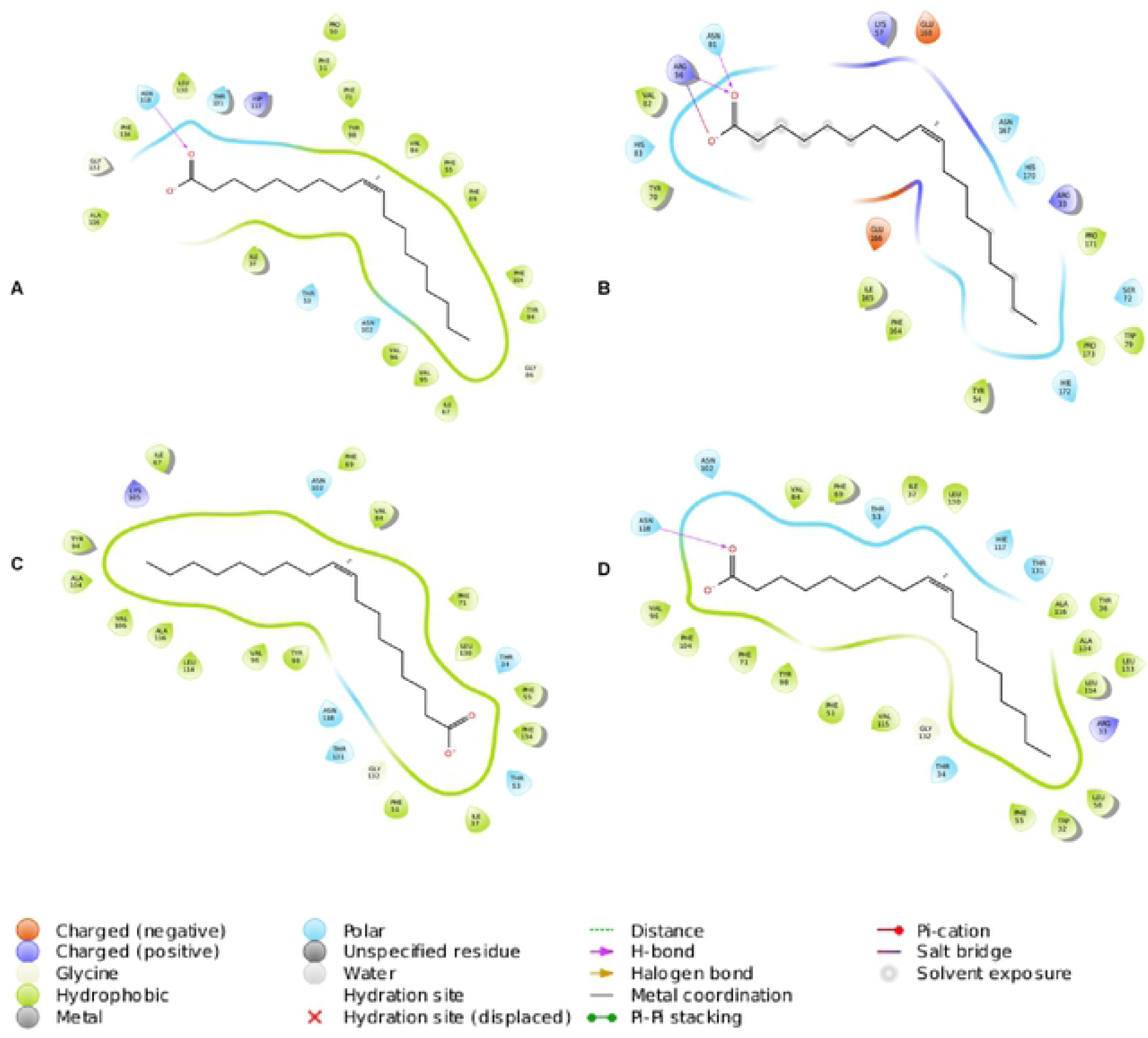
Protein-ligand interactions with oleic acid as ligand docked with OBP1a (A) in pocket 1, OBP1a (B) in pocket 2, OBP1a F104A in pocket 1 (C) and OBP1a F134A in pocket 1 (D)

### *In-silico* predictions on binding of OBP1a and mutants with p-cresol and oleic acid

A systematic approach to prediction of binding of protein and ligand was undertaken using methods of protein-ligand docking, MM-GBSA based affinity determination, molecular dynamic simulations and related analysis, both in the classical ligand binding site as well as in the lateral binding site. These are detailed within Supplementary Text and spans Supplementary Figures. It is observed that, whereas hydrophobic residues contribute to ligands binding in OBP1a, there are compensatory interactions involving charged residues (such as Thr34, Arg52, Arg75 and Lys154) in the four mutants (Supplementary text and Supplementary Figures for details). However, in the case of few mutants, ligand binding is not strong or gets disrupted during the course of molecular dynamics simulations.

At pocket 2, OBP1a WT with oleic acid (−51 kcal/mol) and OBP1a N118A bound to oleic acid (−39 kcal/mol) at pocket 2 show a decreasing trend of binding affinity by 100 ns **(Supplementary Figure 2, Supplementary File 4, Movie Files M4 and M16)** of molecular dynamic simulation of protein-ligand complex. Average binding affinity taken across simulation runs of 100 ns indicate that oleic acid has better binding affinity than p-cresol across both central and lateral binding pockets (**Figure 13; Supplementary File 4**). Taken in context with molecular dynamic simulation results for OBP1a wild type with ligand at pocket1 (**Figures 6 and 7; Movie files M1, M3**) and at pocket 2 **(Figures S1 and S2; Movie files M2 and M4)**, while oleic acid interacts with residues at pocket1 (Gly132, His117) and at pocket 2(Arg56) with high contact strength, p-cresol interacts with residues at pocket 1 including Thr53 thorough stable water bridges and residues like **Phe69, F104, F134** through hydrophobic interactions. Interactions are limited at pocket 2 due to the geometry of the ligand as well as pocket and the biochemical properties of the binding site residues. Although, interactions with oleic acid and OBP1a show higher predicted binding affinity, the highly non-polar enclosed nature of the central cavity would keep p-cresol buried within pocket1 with a slightly higher affinity.

**Figure 13:**
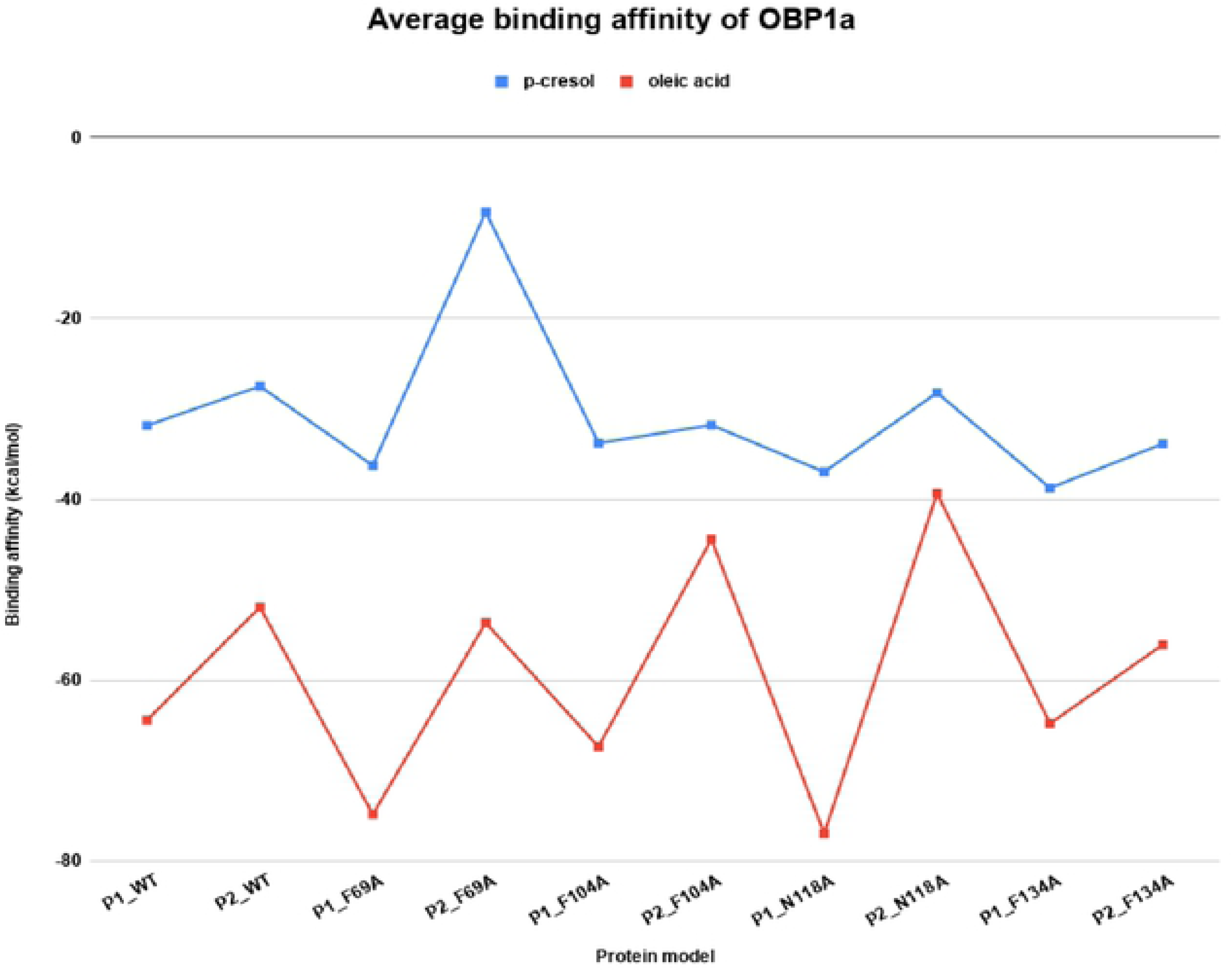
**Thermodynamic binding affinity between OBP1 (WT/mutant) model and ligand (p-cresol/oleic acid) averaged across each 100 ns molecular dynamics simulation trajectory.** P1 is pocket 1 (central cavity) and P2 is pocket 2 (lateral cavi ty). Protein complexed with p-cresol is denoted by blue and that with oleic acid is denoted by red.

**OBP1a F69A** bound to p-cresol at lateral cavity (pocket2) and **OBP1a N118A** bound to p-cresol at pocket2 have poor binding affinities (0.05 kcal/mol and −16 kcal/mol) by 100 ns **(Supplementary Figure 1 and File 4; Movie files M6 and M8)** of molecular dynamic simulation of protein-ligand complex. At 100 ns, in the central cavity (pocket 1), both **OBP1a F134A (Movie file M11) and OBP1a F69A (Movie file M5)** bind to p-cresol separately exhibit a higher binding affinity (−38 kcal/mol and −36.7 kcal/mol) than the wildtype (−35.9 kcal/mol) at either cavities indicating additional stabilizing interactions due to respective mutations. OBP1a bound to p-cresol at pocket 1 (−35.9 kcal/mol) **(Movie File M1)** is more stable than OBP1a bound to p-cresol at pocket 2 (−30 kcal/mol) **(Movie File M2)**. OBP1a F104A at pocket 1 (−33 kcal/mol) **(Movie File M9)**, at pocket 2 (−33 kcal/mol) **(Movie File M10)** and OBP1a F134A at pocket 2 (−34 kcal/mol) **(Movie File M12)** have affinity values in the intermediate range.

In case of mutant F69A, binding seems to be disrupted at pocket 1 as indicated by average binding affinity value but binding at pocket 1 is supported by prolonged hydrogen bonding between Phe51 and p-cresol (**Figure S3**). However, overall, OBP1a F69A binding with p-cresol might not be favourable **(Movie Files M5, M6)**.

The average binding affinity for OBP1a models and oleic acid has a higher range than that of p-cresol. Interestingly, OBP1a N118A binding to each ligand separately at pocket 2 yields similar binding affinity values (**Figure 13**). The complex formed with oleic acid is (−39 kcal/mol), however, highly destabilizing than its wild type (−64 kcal/mol; **Supplementary File 4**). However, this is compensated by the binding at the classical central pocket 1 by OBP1a N118A and oleic acid (−77 kcal/mol). This suggests that OBP1a N118A is likely to bind favourably to oleic acid **(Movie files M15, M16)**.

### Fluorescence binding assays of OBP1a to the sex pheromones

To determine the binding properties of OBP1a, we utilized N-phenyl-1-naphthylamine (NPN) for the competitive fluorescence assay. These binding assays revealed that p-cresol strongly to OBP1a with than oleic acid. This can be attributed to the fact that p-cresol has strong affinity to both central and allosteric pockets whereas oleic acid predicted to bind strongly to the allosteric pocket alone (**S13 D**). We first quantified the ability of OBP1a to reversibly bind NPN. After titration of OBP1a with increasing concentrations of NPN, allowed the measurement of dissociation constant value of 0.7 μM was respectively (**Fig. 14 A**), The wild type OBP1a bind to the fluorescent probe with high yields and strong binding affinity (**Fig. 3A**). The binding affinities of OBP1a with two sex pheromone components, p-cresol and oleic acid were evaluated in competitive binding assays to displace NPN (**Figure. 14 B, C**). OBP1a exhibited strong binding affinities for the two pheromone components, With K_i_ values of 3.5 and 4.2 μM, respectively. These results suggested that, OBP1a may be significant odorant binding protein in male buffaloes, due to its higher binding affinities towards female sex pheromones are likely to be involved in olfactory chemoreception.

**Figure 14:**
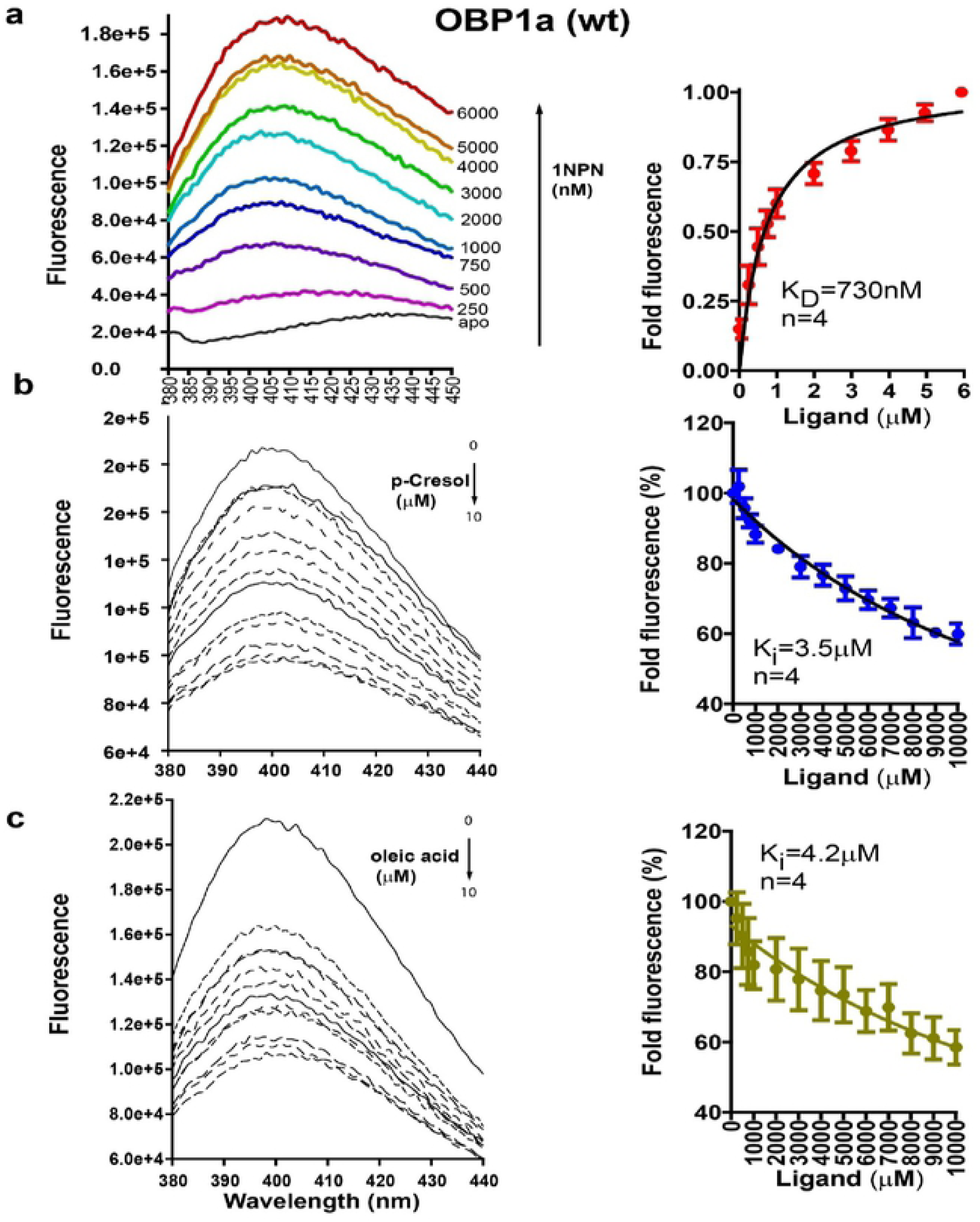
Binding properties of wild type 1OBPa. (A) 1OBPa binds the fluorescent probe 1-NPN with good affinities. (B) and (C)Toward the two sex pheromones (p-cresol and oleic acid) and both exhibited markedly different and complementary spectra of binding.

### Competitive binding assay of ligands with mutants

In order to verify, we used site-directed mutagenesis to evaluate binding cavity and key residues of OBP1a that bind with p-cresol and oleic acid. Phe69, Phe104, Phe134 and Asn118 were directly mutated by alanine and verified by sequencing. Thereafter, competitive binding assay were performed for four mutants OBPa1, with p-cresol and oleic acid. Our results demonstrate that, all mutants did not affect protein solubility and resulted reduced binding affinity with NPN suggesting that other native residues involved ligand-binding process. The recombinant mutant forms (F69A, F104A, F134A and N118A) were used to investigate the binding affinities to the NPN. The binding constants (K_d_) of with 1-NPN were calculated for F69A, F104A, F134A and N118A, as 6 μM, 12.5 μM, 7 μM, and 2 μM, respectively **(Fig. 15 A).** In comparison to the wild-type proteins, four mutants F69A, F104A, F134A and N118A were strongly reduced binding affinities towards p-cresol and oleic acid (**Table 6**). The binding affinity of N118A mutant of isoform OBP1a with oleic acid was higher than both OBP1a-oleic acid and OBP1a-p-cresol, as predicted by MMGBSA results **(Table 4).** As observed from MD simulation analysis, mutation of residues **F69, F104 and F134** disrupts hydrophobic contacts and salt bridges made in the course of protein-ligand binding within the first 100ns itself thus explaining the poor binding of mutants F69A, F104A and F134A. For p-cresol, the mutant F69A lost almost all of its efficient binding capacity whereas, co-mutant N118A showed slightly lower affinity. However, other mutants F104 and F134 have shown non-affinity to p-cresol. Regarding oleic acid, other two mutants F104 and F134A showed weak binding capacity while, F69A and N118 mutants showed slightly lower affinities (**Table 6**). Thus, different site-directed mutagenesis results indicated, none of these compounds could not compete with the 1-NPN even when the concentration of ligand reached in high **(Fig. 15).**

**Figure 15:**
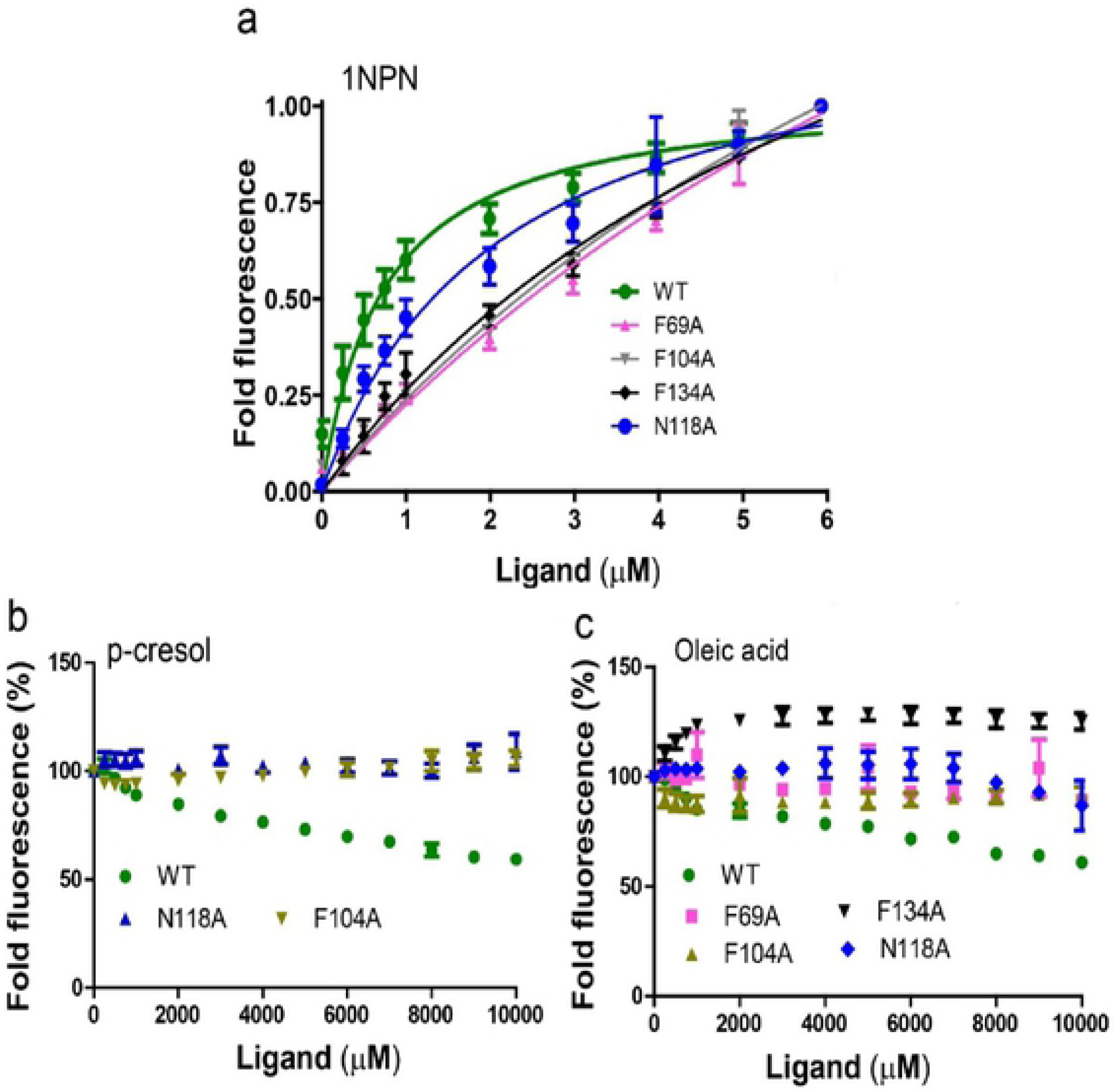
A, B, and C. Competitive binding curves of mutants 1OBPa towards fluorescent probe 1-NPN, and two sex pheromones.

**Table 6:** Binding affinities of buffalo sex pheromones to wild type 1OBPa with four mutants.

| S. No | Proteins | p-Cresol<br>K<sub>i</sub> (μM) | Oleic<br>acid<br>K<sub>i</sub> (μM) |
| --- | --- | --- | --- |
| 1. | WT OBP1a | 3.5 | 4.2 |
| 2. | F69A | ----- | 110 |
| 3. | F104A | ----- | 407 |
| 4. | F134A | ----- | 196 |
| 5. | N118A | 266 | 2.7 |

Note: “----“denotes the protein did not bind with respective ligand in the assay.

## Discussion

Animals utilize various sensory modalities, including olfaction, to communicate to their environment for food, coitus, territorial/dominance relationships and other classical events with both conspecifics [47–50]. In an olfactory event, mechanistically, odorant molecules are solubilized by odorant binding proteins (OBPs) for transport in order to activate odorant receptor (OR) across an aqueous environment in the animal olfactory epithelium [51, 52, 9]. OBPs have been found primarily in olfactory tissues for directly interacting with odorants [53–56, 3, 57] and a few OBPs are expressed in certain body fluids to regulate pheromone-specific behavior [58–60, 20, 61].

Among various well-studied pheromones, p-cresol and oleic acid are known to play an important role in the estrus cycle of the female buffalo. These volatiles when perceived by the male buffalo via the nasal lymph help in regulating mating cues [62, 28]. In this context, we examined the protein expression, localization, binding characteristics and the site-directed mutagenesis of key residues in typical bnOBPs from the buffalo nasal epithelium, the results of which are consistent with a protein that functions in the recognition of female buffalo sex pheromones.

In this study, we built the 3D-structural model for buffalo nasal epithelium odorant-binding proteins (bnOBPs) using the bovine OBP (PDB: 1OBP) structure as template based on their close evolutionary relationship. The predicted models shows typically conserved β-barrel structures with a characteristic central ligand binding pockets with favourable feature of OBP and ligand binding in nature. In mammals, OBPs mainly segregate in nasal epithelial cells and abundant in nasal mucus [53, 63–65, 6]. The tertiary structure of bnOBPs comprises eight β-strands, organized in an anti-parallel β-barrel on one side with α-helices at both terminals. The analysed results, showed 15 major binding sites, preferably for accommodating multiple odorants [28]. Furthermore, the bnOBP isoform models have been validated and nearly 90% amino acid residues were found to be positioned around the favoured regions.

A previous study in our laboratory demonstrated that the proteomics level of buffalo OBPs highly represented in both male and female olfactory glands, the ligand binding and physiology of OBPs remain largely unclear [26, 28]. We report here that *in-silico* and fluorescence binding studies comparison of wild type and mutant forms indicates specific role of OBP1a in the detection of sex pheromones. Since, OBP1a is predicted to have distinct binding properties compared to other two OBPs, we used site-directed mutagenesis and fluorescence quenching assays to characterize binding abilities of the four mutants of OBP1a. Our result reveals high affinities of bnOBPs with sex pheromones, in particular, the interaction between OBP1a and tested ligands indicating that OBP1a may be involved in sex pheromone recognition and transportation. Interestingly, OBP1d and OBP1f proteins were predicted to have weak interactions with the sex pheromones, implying that those proteins may not be the main protein that responds to sex pheromones and that it may have different functions.

From a BLAST research in the PDB, bovine OBP (PDB: 1OBP) with most sequence similarity (90% identity) to tabbed as the suitable template to build a 3D homology structure of bnOBPs. The tertiary structure of bnOBPs comprises eight β-strands, organized in an anti-parallel β-barrel on one side with α-helices at both terminals. Interestingly the deduced amino acid sequences suggested that, the most important structure, beta (TIM) barrel, is depicted in bnOBPs, which would provide a favourable binding position to the odor/chemical cues.

We found that among the OBP isoforms isolated from the nasal lymph, isoform **OBP1a** is the most identical to its phylogenetic relative, bovine OBP. We have found two strong candidates for ligand-binding sites on OBP1a isoform, one centrally located in the beta barrel and the other laterally located between the beta barrel and near the C-terminal alpha helix/loop region. Docking scores predicted better poses at the central cavity (pocket 1) for both ligands whereas thermodynamic affinities predicted suggested that oleic acid is more likely to bind at the allosteric pocket. Oleic acid being longer chained with an absence of an aromatic ring might indeed be more likely to bind to a more easily accessible binding site. On the other hand, small aromatic ligands like p-cresol could bind to both sites but would be highly favoured at a core hydrophobic cavity in the beta barrel of the OBP1a.

Based on protein-ligand interactions observed via docking, homology models for mutants were constructed to understand the energetics and dynamics of protein-ligand binding. Molecular dynamic simulations uncovered that p-cresol binds to the central cavity in isoform OBP1a predominantly through an alternating interplay of hydrogen bonding and water bridge contacts among certain residues. The mutant models in complex with p-cresol show loss of hydrogen-bonding and remarkably negligible water-mediated and hydrophobic contacts throughout the simulation suggesting that OBP1a and p-cresol would bind strongly whereas mutations F69A and N118A would disrupt the equilibration of the complex result in poor/ineffective binding. However, site at pocket 2 or sites newly exposed in a given mutant model during the course of ligand-binding might account for binding affinities in the weaker range.

OBP1a complexed with oleic acid and OBP1a F104A-oleic acid at the central cavity shows a partial disruption in helicity near the N-terminus. Ionic bonds are indicated in ligand interactions with F104A and F134A models although OBP1a F134A does not equilibrate within 100ns. Overall, changes in helicity and heavy atom fluctuations as well as dynamic residue-ligand contacts suggest that while oleic acid would bind strongly to the central cavity of OBP1a, mutant models F104A and F134A would bind poorly to oleic acid due to loss of water-mediated contacts and time taken to achieve equilibration respectively. Our bioinformatics results are in agreement with experimentally observed results for binding of ligands p-cresol and oleic acid each to isoform OBP1a. The presence of salt bridges, hydrophobic contacts and hydrogen bonds helps stabilize protein-ligand binding in case of mammalian odorant binding, OBP1a. Other isoforms of OBP1 are also likely to show similar binding profiles with estrus-specific pheromones p-cresol and oleic acid due to high sequence and structural homology with OBP1a. In this scenario, presence of isoforms with similar binding properties would compensate for loss in an isoform of OBP1, thus maintaining robustness in binding function. Based on our results and in addition to mutations F69A and N118A, it would be worthwhile to investigate experimentally mutating Phe51 and Thr53 in OBP1a to test binding with p-cresol. In case of oleic acid as ligand and in addition to mutations F104A and F134A, residues His117 and Gly132 should also be investigated experimentally due their distinct roles in ligand interactions as per our results.

## Conclusion and Future perspective

A systematic bioinformatics approach to identify protein-ligand binding in the case of mammalian odorant binding protein (OBP) has helped us identify the extent of homology, putative binding sites, likelihood of the ligand in complex with the protein and dynamics of the protein-ligand complex for 100 ns in an aqueous simulation box mimicking the environment in the nasal mucus covering the olfactory epithelium of the male buffalo. In particular, the site-directed mutagenesis strategy has led us to understand the residues in OBP1a during recognition of sex pheromones.

This is the first experimental report that the six different OBP isoforms from the buffalo nasal epithelium were characterised by molecular, biochemical and structural approaches. The binding affinity and other localization diversity of OBPs are useful for understanding the physiological role and mechanism of olfactory communication in the buffalo. Notably, fluorescence competitive studies together with mutational studies furnish empirical evidence for the *in-silico* predicted functions of specific amino acids in OBP1a. We speculate that our study would further play a crucial guiding role in the future development of a biosensor for estrus detection in buffalo.

## Materials and Methods

### RNA extraction and cDNA conversion

Fresh nasal epithelial tissue sample has been isolated within 2 hrs from sacrificed male buffalo from a slaughterhouse. The tissue samples were washed with 1X PBS and flash frozen in liquid nitrogen and stored at –80° C. Total mRNA was extracted from the nasal epithelium of buffalo with the Trizol Reagent kit (Invitrogen™ TRIzol™ Reagent, USA) as per manufacturer protocol. The quantity and integrity of RNA was measured using Nano Drop 2000c spectrophotometer (Thermo Scientific, USA). Using freshly isolated RNA, the First-strand cDNA was synthesized using RevertAid First Strand cDNA Synthesis Kit (Thermo Scientific, USA) and employed as templates for further gene amplification.

### Molecular cloning and expression vectors construction

The gene-specific primers (Forward 5’—ATATA<u>CATATG</u>AAGGTTCTGTTCCTGACTC—3’ & reverse 5’—TATAT<u>GTCGA</u>CTCACTCGGGGTGAGGATG—3’) were designed using *Bubalus bubalis* OBP sequence (GenBank accession: 006042607) as a reference. These primers are having recognition sites for *NdeI* and *Sal*I for directional cloning into the expression vector. Using first strand cDNA (1 μL) as a template, OBP gene was amplified in a ProFlex™ PCR System (Applied Biosystems, USA) in a total reaction solution (25 μl) using Q5 High-Fidelity DNA polymerase (NEB) with 1 μM of each PCR primer. The PCR cycle were as followed: After a first denaturation step at 98 °C for 1 min, 35 amplification cycles (10 sec at 98 °C, 10 sec at 60°C, 15 sec at 72 °C) were performed, followed by a final step of 2 min at 72 °C. The PCR product, with the expected 522 bp size was purified from the agarose gel using Wizard® SV Gel and PCR Clean-Up System (Promega, USA) according to the manufacturer’s instructions. The purified PCR product was then ligated into the plasmid PCR®II-TOPO® using the TOPO TA cloning kit (Invitrogen, USA). After transformation, positive clones were confirmed by blue white screening as well as gene specific colony PCR using above mentioned primers. From putative positive clones, the recombinant plasmids were isolated using The Wizard® Plus SV Minipreps DNA Purification System (Promega, USA). The plasmids having OBP insert DNA was sequenced by Illumina sequencing (NCBS, India). The positive Topo/OBP plasmid DNA containing the appropriate OBPs sequences was digested with *Nde*1 and *Sal*1 restriction enzymes for 4 hrs at 37 °C and purified using Wizard® SV Gel and PCR Clean-Up System (Promega, USA) as per the manufacturer’s instructions. Similarly, bacterial expression vector pET28a+ was also linearized with the same enzymes and the desired linearized DNA was also purified. The digested OBP PCR product and pET28a+ vector were ligated using T4-DNA ligase for 16 hr at 18 °C and followed by transformation to ultra-competent *E*. *coli* XL10-Gold® cells (Agilent, USA). The positive clones were screened using colony PCR and further sequenced. We have isolated six isoforms of bnOBPs. The computation of various physical and chemical parameters for the bnOBPs was done using the online tool ProtParam on the Expasy SIB Bioinformatics Resource Portal (http://expasy.org/) [66].

### 3D model construction and molecular docking

#### Template selection

A BLAST search was used [67] to find a suitable sequence alignment for three dimensional structure construction. The amino acid sequence of bnOBPs was used to find the corresponding structural templates in PDB (http://www.rcsb.org) database. The crystalline structure of odorant-binding protein from bovine nasal mucosa of *Bos taurus* (PDB ID: 1OBP, Chain A, resolution 2.0 Å) matched with 90% identity with isolated bnOBPs sequences and was hence further selected as a template.

#### Structural modelling

The sequence alignment between target and the template was conducted by using Clustal omega software [68]. The PEP-FOLD server [69, 70], which builds on a new *de novo* approach to predict 3D peptide structures from sequence information for lacking amino acid residues from template. The aligned sequence was structurally modelled using MODELLER software (v9.20) modelling [71]. A set of 20 models were generated from MODELLER, optimum alignment was determined by the lowest QMEAN4 score and discrete optimized energy (DOPE) scores. The modelling rationality was further estimated using Pro-CHECK (http://nihserver.mbi.ucla.edu/SAVS/) [72]. The folding pattern and homogeneity comparison was predicted using PyMol software (PyMOL Graphics software, v2.3, <u>Schrödinger</u>). Structural superimposition was carried out for constructed models with corresponding template [28].

### Homology with bovine OBP

Blastp [73] was performed with the subject sequence as bovine OBP (PDB ID: 1OBP, Chain A) and query sequences OBP1a, 1b, 1c, 1d 1e and 1f. BLOSUM62 matrix was used with a word size of 3 and expect threshold of 10.

### Multiple Sequence Alignment of bnOPB isoforms

Protein isoforms were aligned using MAFFT (v7.452) [74] with global alignment with an FFT approximation (G-INS-i) and a maximum of 1000 iterations. The resultant sequence alignment in CLUSTAL format was visualized in ESPript 3.0 [75] to generate alignment outputs. Phylogeny of bnOBPs isoform was generated using the Neighbour-Joining method in ClustalW2 [76] and visualized in iTOL [77]. The tree was rooted to bovine OBP sequence (PDB: 1OBP_A; [16]).

### Domain analysis

Isoform models were evaluated for the presence of lipocalin domain using HMMSCAN [78] with Pfam database, v32.0.

### Protein preparation

Each of the models was prepared using the Protein Preparation Wizard [79] in MAESTRO version 11.9.011 [80–86] on Linux platform, Schrodinger Inc. Each model was corrected for bond orders, bond types, missing atoms and charge keeping the pH between 6.4 to 7.2, in order to mimic the natural chemical environment in the buffalo nasal lymph. Waters beyond 5 Å were deleted from heteroatom groups. Water molecules with less than three hydrogen bonds to non-water molecules were removed. Protonation and orientation states were corrected accordingly and the hydrogen bond network was optimized to aid protein preparation following which restrained minimization was carried out to allow hydrogen atoms to be minimized using OPLS3e [87, 88] force field and to relax strained bonds, angles and clashes.

### Site Analysis

Each minimized protein model corresponding to each of the 11 isoforms was further tested for potential ligand-binding and druggable sites that were identified using a restrictive definition of hydrophobicity and a standard grid with crop site maps at 4 Angstrom cut off from site point in Sitemap [89–91]. This was done to identify top-ranked potential receptor-binding sites. The ligand-binding score (Site Score) and druggability (D-score) were used for filtering predicted binding sites with a cut-off of 0.8 each. A score greater than or equal to 1 indicates a strong probability of being a potential binding site whereas a score between 0.8 and 1 indicates a partial binding site. The site corresponding to a **central cavity (pocket 1)** was chosen further for preparation of receptor grid and docking against ligand. A receptor grid was generated corresponding to each potential binding site using Van der Waals radius scaling factor of 1 and a partial charge cut-off of 0.25. The docked ligand was confined to the enclosing box and the centroid was taken into account. The force field used was OPLS3e.

### Ligand selection and preparation

Ligands were chosen based on their relevance as potential mating cues and status as estrus-specific pheromone compounds released during the estrus phase of the female buffalo [50]. A ligand library was prepared comprising pheromones p-cresol (CID: 2879) and oleic acid (CID: 445639) and fluorophore 1-NPN (CID: 7013) to test binding (**Table 7**). The two-dimensional information on compounds was obtained in SMILES [92] format from Pubchem [93]. Force field OPLS3e was used to generate possible ligand states in the pH range 6.4 to 7.2 including tautomers using Epik [94, 95]. All combinations of chiralities were generated for the target pH interval. Accurate, energy-minimized three-dimensional molecular structures of ligands were generated using LigPrep [91]. The ligands were desalted and tautomers were generated. Chiralities were determined from 3D-structure. A final ligand library was prepared which was docked against each potential binding site of the docked model.

**Table 7:**
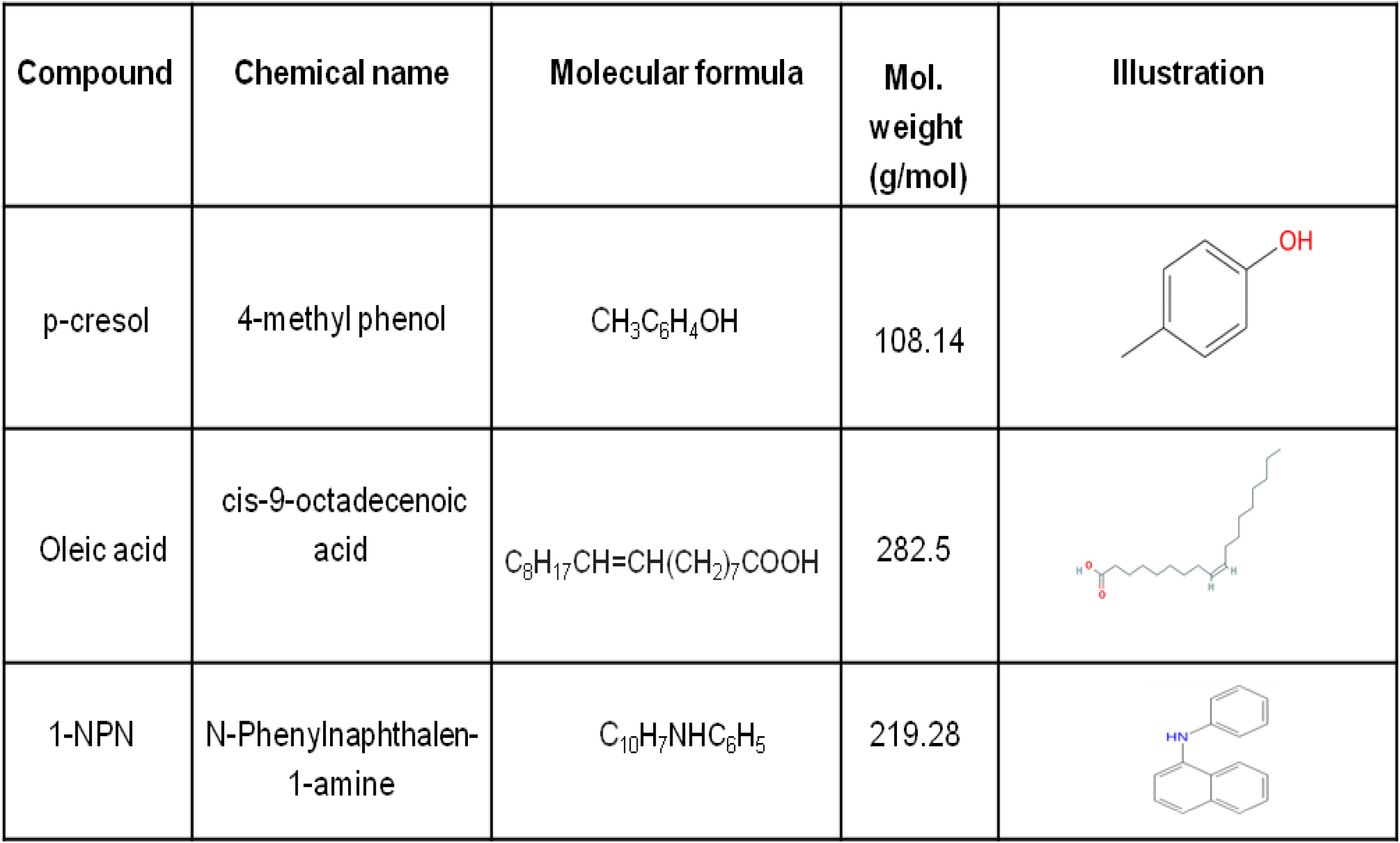
Ligand structures used for docking A. p-cresol B. oleic acid and C. 1-NPN

| Compound | Chemical name | Molecular formula | Mol. weight (g/mol) | Illustration |
| --- | --- | --- | --- | --- |
| p-cresol | 4-methyl phenol | $\text{CH}_3\text{C}_6\text{H}_4\text{OH}$ | 108.14 | |
| Oleic acid | cis-9-octadecenoic acid | $\text{C}_8\text{H}_{17}\text{CH}=\text{CH}(\text{CH}_2)_7\text{COOH}$ | 282.5 | |
| 1-NPN | N-Phenylnaphthalen-1-amine | $\text{C}_{10}\text{H}_7\text{NHC}_6\text{H}_5$ | 219.28 | |

### Protein-ligand docking for wild type and mutant isoforms

Protein-ligand docking was carried out with Glide (96, 97, 89, 90] in the presence of ligands, oleic-acid, p-cresol and 1-NPN. Ligands with less than or equal to 500 atoms and 100 rotatable bonds were considered and scored. To soften the potential for non-polar parts of the ligand, the van der Waals radii of ligand atoms was scaled with partial atomic charge less than the specified cut off with a scaling factor of 0.80 and a partial charge cut off of 0.15.

### Protein-ligand binding affinity for wild type and mutant isoforms

The minimized model of each isoform was tested against this ligand library set for predicting relative binding affinity using the MM-GBSA protocol in Prime [98, 99]. The solvation model used was VSGB with the OPLS3e force field with sampling method as minimization of whole complex. Thermodynamic binding affinity was calculated between ligands and models generated for OBP (wild type and mutants) isoforms and ligands using MM-GBSA in Maestro Prime, Schrodinger LLC. This approach is used to predict the free energy of binding for set of ligands to receptor. The docked poses were minimized using the local optimization feature in Prime and the ligand strain energies. Energies of the ligand-receptor complexes were calculated using Prime MM-GBSA technology with all receptor residues being held frozen.

The binding free energy ΔG_bind_ is estimated using following equation

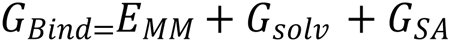

ΔE_MM_ = difference in energy between the complex structure and the sum of the energies of the ligand and unliganded protein using the OPLS3e force field.

ΔG_solv_ = difference in the GBSA solvation energy of the complex and the sum of the solvation energies for the ligand and unliganded protein.

*Δ*G_SA_ = difference in the surface area energy for the complex and the sum of the surface area energies for the ligand and unliganded protein.

### Molecular Dynamic simulations for wild type and mutant OBP1a with ligands

Molecular Dynamic simulation of OBP1a wild type and virtual mutants were performed in the absence of ligand and in the presence of p-cresol/ oleic acid for a simulation time of 100 ns each using DESMOND [100]. Molecular dynamic (MD) simulations were carried out on docked models of buffalo OBP wild type (WT) isoform with each ligand (p-cresol and oleic acid) for a time period of 100 nanoseconds. The system was built using the TIP4P water model with an orthorhombic box shape of dimensions 10 Å uniformly. The force field used was OPLS3e. The system was neutralized by adding appropriate ions and recalculated. In order to mimic the chemical environment in the nasal mucus of the buffalo, sodium chloride (NaCl) of concentration 0.15 M was used. The system was built using TIP4P water model with an orthorhombic box shape of 10 Å distance from xyz axes. The volume was minimized and the force field used was OPLS3e. The simulation time was 100 ns with temperature 300K and pressure 1.01325 bar. The model system was relaxed before simulation. The thermostat used was Nose-Hoover chain and barostat used was Martyna-Tobias-Klein method with an isotropic coupling style. Interactions chosen were Coulombic with a short-range cut-off method and cut-off radius of 9Å. Velocities were randomized for the simulation. The system was appropriately relaxed before the production run. Simulation analysis of each system was performed with respect to DSSP, RMSF, RMSD, H-bonds and energy. The criteria for hydrogen bonds is also in addition to the presence of atoms, the right geometry criteria which is distance <=2.5 Å, donor angle is >=120 degrees and acceptor angle is >=90 degrees. Pi-pi stacking interactions were also analysed within 4.5 Å in addition to hydrophobic interactions between a side chain and aromatic/aliphatic carbons of the ligand within a 3.6 Å cut-off.

### Thermal stability of MD snapshots

Snapshots from each 1000 frame-MD trajectory were extracted at an interval of 100 frames or 10 ns. MM-GBSA score was assigned for each snapshot comprising protein and ligand co-ordinates. Python script *thermal_mmgbsa.py* from Schrodinger and an in-house custom Python script were used for this purpose. The average binding affinity for a given protein-ligand complex was calculated by taking the mean of the thermal binding affinity across the ten frames extracted from each such trajectory.

### Recombinant protein expression and purification

To express and purify the 6x Histidine tagged bnOBP recombinant proteins (6X-His-bnOBP), the pET28a+OBP1a recombinant plasmid were transformed into Rosetta™(DE3) Competent Cells (Novagen). The positive clones were inoculated in 10 mL LB medium having kanamycin (50 μg/mL) under constant shaking at 200 rpm at 37 °C. After 6 hrs, the culture was inoculated to 1 litre of LB medium and grown till OD (600nm) of 0.5 to 0.7. To induce protein expression, 0.5 M Isopropyl-β-D-thiogalactopyranoside (IPTG) 1 ml was added and culture was further grown overnight at 16 °C under constant shaking. Cells were harvested using centrifugation at 7000 x g for 30 min and cell pellet was washed with 50 mM Trish-HCl pH 7.6. Further the cell pellet was resuspended in lysis buffer (50 mM Trish-HCl, 300 mM NaCl, 1% Triton X-100, 5 mM beta-Mercaptoethanol, 50 mM Imidazole, pH 7.6).The suspension was sonicated for 7 cycles of 5 seconds pulse with 3.30 mins intervals at 40% of the maximal acoustic power using a sonicator on ice (Sonics & Materials, Inc. USA). The sonicated cell lysate were further centrifuged at 35000 x g for 45 min yielded a clear supernatant called as cell free extract containing the soluble 6X-His-bnOBP protein. The histidine tagged protein was purified using Ni-NTA affinity chromatography (Invitrogen, USA) according to the manufacturer’s protocol. In brief, it is equilibrated the column with 10 ml of wash buffer (50 mM Tris Hcl, 300 mM NaCl, 50 mM imidazole, and 2 mM beta-Mercaptoethanol, pH 8.0), and added the sample. Then, the column was washed with 10 ml of binding buffer and eluted the column with 2 ml of elution buffer (50 mM Tris Hcl, 300 mM NaCl, 100mM – 500mM imidazole, 2mM beta-Mercaptoethanol, pH 8.0) and collected the eluates into four separate 1.5 ml Eppendorf tubes in chronological order 2 ml per tube. The size and purity of recombinant bnOBPs were confirmed by 12% Sodium dodecyl sulphate polyacrylamide gel electrophoresis (SDS-PAGE) analysis. The concentration of the purified proteins was measured using BCA protein assay kit (Thermo Scientific, USA).

### Binding assays

Fluorescence binding assays were performed on a Horiba Scientific Fluoromax-4 spectrofluorometer using slits of 3, 4, or 5 nm according to the protein and a light path of 1 cm. The OBP1a were dissolved with 50 mM Tris·HCl, 300mM Nacl (pH 8.0) at a final concentration of 2 μM. The N-phenyl-1-naphthylamine (1-NPN) fluorescent probe was used as a fluorescent reporter at 2 μM concentration and titrated with each competitive ligand at between 0.1 and 13 μM concentrations. The sample was excited at 337 nm, and emission was collected from 380 and 450 nm. The affinity of two ligands was measured in competitive binding assays. Dissociation constants for 1-NPN were calculated using the Graph Pad Prism software. Fluorescence measurements were performed according to Zhu *et al* 2017 [21]. Dissociation constants of the competitors were calculated by the equation,

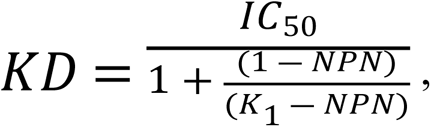

Where *IC*_50_ is the concentration of ligands with the initial fluorescence value of 1-NPN, [1-NPN] is the free concentration of 1-NPN, and K1-NPN is the dissociation constant of the complex protein/1-NPN. All values reported were performed in triplicates, except for ligands showing not significant binding that were analysed in single experiments.

### Simulation of Site-directed mutagenesis and the expression of mutants

According to the molecular docking studies, OBP1a was choosed to further evaluation. We predicted that four key binding sites in the process of OBP1a binding with tested ligands. For site-directed mutagenesis, we used OBP1a construct as template, designed and prepared four mutants of this OBP1a by replacing mutations of, F69A (mutating phenylalanine to alanine at position 69), F104A (mutating phenylalanine to alanine at position 104), P134A (mutating phenylalanine to alanine at position 134) and N118A (mutating asparagine to alanine at position 118) were generated by PCR using specific mutated primers and also listed in **Table 8** The mutants were verified by plasmid DNA sequencing. The OBP1a recombinant mutant proteins prokaryotic expression and purification were conducted as described above and used for competitive binding experiments.

**Table 8.**
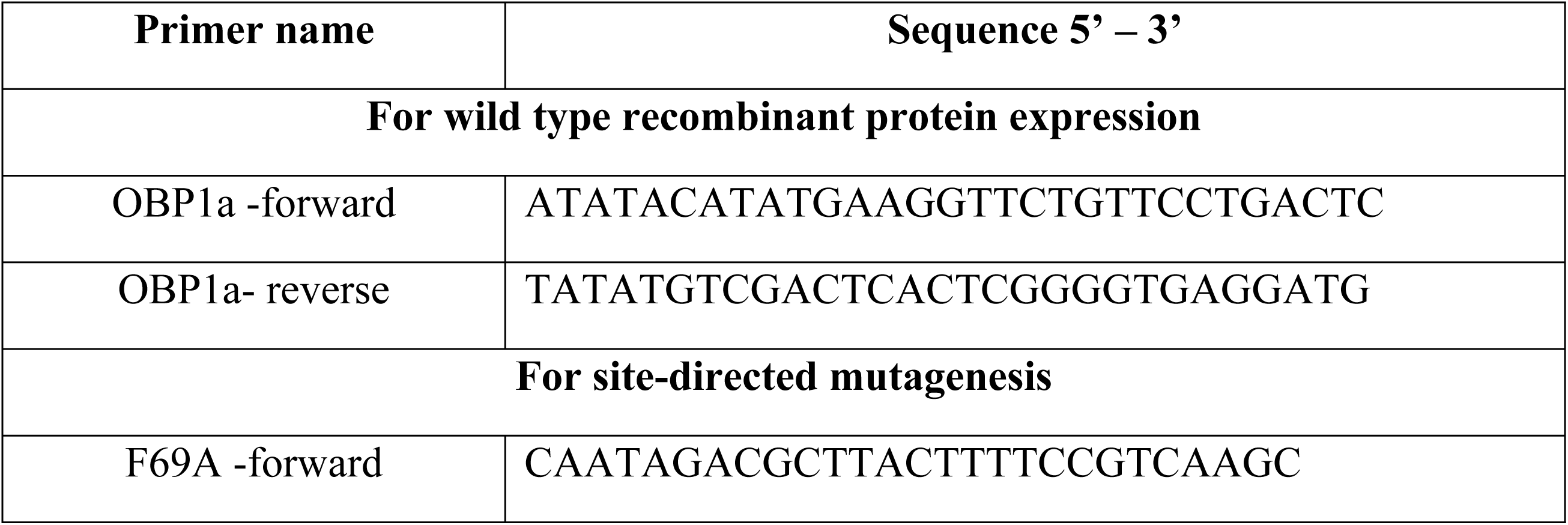
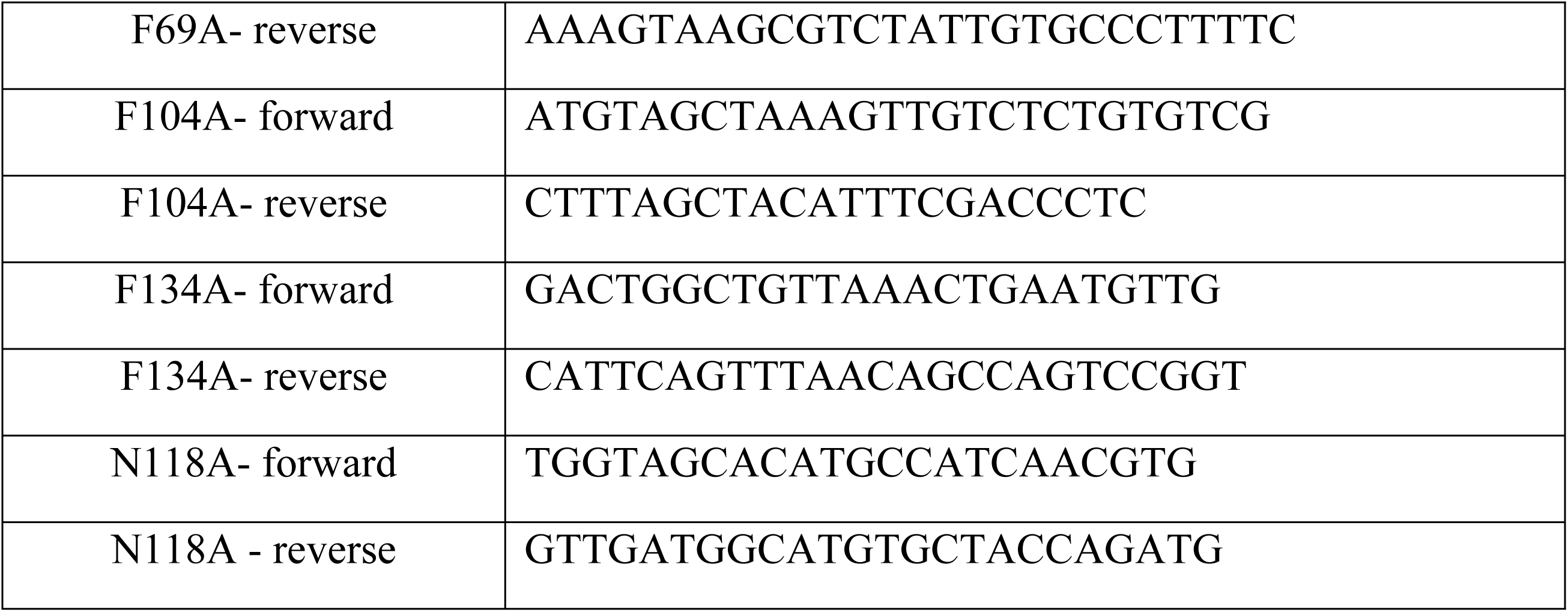
List of primers used for site-directed mutagenesis.

## Supporting information

**Supplementary file1:** bnOBPs, Table: residue interactions

**Supplementary File2:** Supplementary Tables

**Supplementary File3:** Supplementary Text

**Supplementary File4:** Supplementary figures

## List of Supplementary Figures

### List of Figures

1. **S1:** Molecular dynamic simulation of OBP1a – p-cresol for 100 ns in lateral cavity (pocket 2)
2. **S2:** Molecular dynamic simulation of OBP1a – oleic acid for 100 ns in lateral cavity (pocket 2)
3. **S3:** Molecular dynamic simulation of OBP1a F69A – p-cresol for 100 ns in central cavity (pocket 1)
4. **S4:** Molecular dynamic simulation of OBP1a F69A – p-cresol for 100 ns in lateral cavity (pocket 2)
5. **S5:** Molecular dynamic simulation of OBP1a N118A – p-cresol for 100 ns in central cavity (pocket 1)
6. **S6:** Molecular dynamic simulation of OBP1a N118A– p-cresol for 100 ns in lateral cavity (pocket 2)
7. **S7:** Molecular dynamic simulation of OBP1a F104A – p-cresol for 100 ns in central cavity (p1)
8. **S8:** Molecular dynamic simulation of OBP1a F104A – p-cresol for 100 ns in lateral cavity (pocket 2)
9. **S9:** Molecular dynamic simulation of OBP1a F134A – p-cresol for 100 ns in central cavity (pocket 1)
10. **S10:** Molecular dynamic simulation of OBP1a F134A – p-cresol for 100 ns in lateral cavity (pocket 2)
11. **S11:** Molecular dynamic simulation of OBP1a F69A – oleic acid for 100 ns in central cavity (pocket 1)
12. **S12:** Molecular dynamic simulation of OBP1a F69A – oleic acid for 100 ns in lateral cavity (pocket 2)
13. **S13:** Molecular dynamic simulation of OBP1a N118A – oleic acid for 100 ns in central cavity (pocket 1)
14. **S14:** Molecular dynamic simulation of OBP1a N118A – oleic acid for 100 ns in lateral cavity (pocket 2)
15. **S15:** Molecular dynamic simulation of OBP1a F104A – oleic acid for 100 ns in central cavity (pocket 1)
16. **S16:** Molecular dynamic simulation of OBP1a F104A – oleic acid for 100 ns in lateral cavity (pocket 2)
17. **S17:** Molecular dynamic simulation of OBP1a F134A - oleic acid for 100 ns in central cavity (pocket 1)
18. **S18:** Molecular dynamic simulation of OBP1a F134A – oleic acid for 100 ns in lateral cavity (pocket 2)

**Movie files** M1 to M20 referenced throughout the manuscript have been provided as supplementary files in the following link.

https://drive.google.com/drive/folders/1oG5PwG3Ub-CKCR0QJkl0lz9Mp0lGKFes?usp=sharing

## Abbreviations

OBP: Odorant binding proteins
OR: odorant receptor
CSP: chemosensory proteins
NPC2: Niemann-Pick type C2 proteins
ORF: open reading frame
RMSD: root mean square deviation
WT: wild type
MD: Molecular dynamics
RMSF: root mean square fluctuations
MM-GBSA: molecular mechanics energies combined with the generalized born and surface area continuum solvation
NPN: N-phenyl-1-naphthylamine
BLAST: basic local alignment search tool
TIM: triosephosphateisomerase
PBS: phosphate buffer saline
PCR: polymerase chain reaction
QMEAN: qualitative model energy analysis
DOPE: discrete optimized energy
BLOSUM: blocks substitution matrix
MAFFT: multiple alignment using fast fourier transform
HMMSCAN: hidden Markov models scan
Pfam: protein families
OPLS: orthogonal projections to latent structures.

## Acknowledgements

The facility availed from DST-FIST and DST-PURSE, Government of India is acknowledged. Dr. GA for award of UGC-BSR Faculty Fellowship (No. F. 18-1/2011(BSR) dt.04.01.2017). Dr. Akash Gulyani and his colleagues acknowledge support at NCBS-inStem. RS would like to acknowledge her JC Bose Fellowship (JC Bose fellowship (SB/S2/JC-071/2015) from Science and Engineering Research Board and NCBS (TIFR) for infrastructural facilities. BM would like to acknowledge Tata Trust-TDU Fellowship for PhD awarded to her from 2017 to 2019.

## Author Contributions

**Conceptualization:** Chidhambaram Manikkaraja, Bhavika Mam, Randhir Singh.

**Data curation:** Chidhambaram Manikkaraja, Bhavika Mam, Randhir Singh, Balasubramanian Nagarathnam, Geen George.

**Formal analysis:** Chidhambaram Manikkaraja, Bhavika Mam, Randhir Singh, Balasubramanian Nagarathnam, Geen George.

**Funding acquisition:** Ramanathan Sowdhamini.

**Investigation:** Chidhambaram Manikkaraja, Bhavika Mam, Randhir Singh.

**Methodology:** Chidhambaram Manikkaraja, Bhavika Mam, Randhir Singh, Balasubramanian Nagarathnam, Geen George.

**Project administration:** Ramanathan Sowdhamini.

**Resources:** Ramanathan Sowdhamini, Akash Gulyani, Govindharaju Archunan.

**Supervision:** Ramanathan Sowdhamini.

**Validation:** Chidhambaram Manikkaraja, Bhavika Mam, Randhir Singh.

**Visualization:** Chidhambaram Manikkaraja, Bhavika Mam, Randhir Singh.

**Writing –** Chidhambaram Manikkaraja, Bhavika Mam, Randhir Singh.

**Writing – review & editing:** Ramanathan Sowdhamini, Govindharaju Archunan, Akash Gulyani.

